# Differential effects of two common GVHD prophylaxis regimens on the gut microbiome: Results from the BMT CTN 1801 study

**DOI:** 10.64898/2026.02.19.706769

**Authors:** Jakob Wirbel, Wael Saber, Michael J. Martens, Jonathan U. Peled, Tessa M. Andermann, Teng Fei, Erin F. Brooks, Boryana Doyle, Nathan B. Pincus, Robert R. Jenq, Merav Bar, Javier Bolaños-Meade, Brandi Bratrude, Saurabh Chhabra, Sung Won Choi, William Clark, Suman Das, Hany Elmariah, Mahasweta Gooptu, Shernan G. Holtan, Richard J. Jones, John E. Levine, Brent R. Logan, Monzr M. Al Malki, Hemant S. Murthy, Armin Rashidi, Andrew R. Rezvani, Marcie L. Riches, Lyndsey Runaas, Karamjeet Sandhu, Ashley Spahn, Anthony D. Sung, Marcel R.M. van den Brink, Mary M. Horowitz, Mehdi Hamadani, Leslie S. Kean, Miguel-Angel Perales, Ami S. Bhatt

## Abstract

Allogeneic hematopoietic cell transplantation (allo-HCT) is a potentially curative treatment for many hematological malignancies, but graft-versus-host disease (GVHD) is a common complication. Low gut microbiome diversity is associated with higher GVHD risk and shorter survival in multiple studies.

Recently, the BMT CTN 1703 clinical trial demonstrated superiority of a GVHD-prophylaxis regimen including post-transplant cyclophosphamide (PTCy) compared to the standard prophylaxis (tacrolimus and methotrexate, Tac/MTX) in terms of GVHD-free, relapse-free survival at one year among reduced intensity conditioning allo-HCT recipients. However, the effect of PTCy on the gut microbiome and its association with clinical outcome have not been described. Here, we report on a companion randomized clinical controlled trial (BMT CTN 1801), which collected 2575 longitudinal stool samples from 304 study participants. Samples were obtained up to weekly up to day 84 post allo-HCT and at less frequent intervals thereafter, up to 2 years.

Microbiome diversity and absolute microbial load were lower in the PTCy group compared to the Tac/MTX group on days 14–28 post-HCT. However, diversity at the timepoint closest to neutrophil engraftment was not significantly associated with non-relapse mortality after one year or other clinical outcomes, contrary to expectations from previous studies. Microbial domination events, when a single species exceeds 30% relative abundance, were comparable across treatment arms and reflected both pathogen blooms as well as less severe disruptions of the microbial community. *Clostridium scindens* and secondary bile acid metabolism pathways were less prevalent in the PTCy arm than in the Tac/MTX arm post-HCT, yet presence of secondary bile acid metabolism pathways was associated with a lower risk of chronic GVHD.

Given that PTCy was associated with a greater disruption of the microbiome as measured by diversity, absolute microbial abundance, and bile acid metabolism capability, but better clinical outcomes overall, these data suggest that the importance of the microbiome in modulating the host immune systems after allo-HCT is specific to different types of GVHD prophylaxis.

## Introduction

Hematologic malignancies were responsible for around 9% of new cancer cases and deaths in the US in 2024 (ref^1^). A key therapeutic approach for hematologic cancers is allogeneic hematopoietic cell transplantation (allo-HCT), consisting of cytotoxic chemotherapy with or without irradiation followed by infusion of hematopoietic cells from a donor matched for some or all of the major histocompatibility alleles. While this treatment can potentially cure the underlying malignancy, a major and common complication of allo-HCT is graft-versus-host disease (GVHD). GVHD occurs when donor T-cells attack the host tissues, resulting in both acute^2^ and chronic^3^ immune-related complications. GVHD is a significant contributor to mortality in individuals undergoing allo-HCT.

Strategies to prevent GVHD include optimizing human leukocyte allele (HLA) matching as well as immunosuppressive regimens, including the standard therapy consisting of methotrexate (MTX) and calcineurin inhibitors^4^. However, GVHD still develops in more than half of the patients treated with this regimen. Several approaches exist to improve GVHD prevention through inhibiting graft T-cells alloreactivity: T-cell depleted grafts^5^, grafts enriched for regulatory T-cells^6^, and ablation of allo-reactive T-cells post-HCT using post-transplant cyclophosphamide (PTCy)^7^. Recently, the BMT CTN 1703 trial reported superior efficacy for PTCy over the standard calcineurin inhibitor/methotrexate regimen in terms of GVHD-free, relapse-free survival at one year in a prospective, multi-center, randomized controlled clinical trial^8^.

Prior studies show the gut microbiome, especially gut microbiome diversity, to be associated with adverse outcomes in allo-HCT, such as risk for GVHD^9^, bloodstream infections^10^, and mortality^11^. For example, microbial domination, where a single taxon accounts for more than 30% of the sequencing reads in a sample, is associated with bacterial bloodstream infections and other poor clinical outcomes^10^. In a large observational study encompassing over 1000 participants from four institutions, overall survival after one year was longer in participants with higher microbial diversity^11^. Interestingly, this association was not observed for T-cell depleted grafts, suggesting that a diverse gut microbiome might be important for dampening the alloreactivity of graft T-cells. As a consequence of these findings, efforts are underway to test fecal microbiota transplantation as a method to ‘rescue’ low microbial diversity and potentially mitigate the adverse effects of GVHD^12^.

The mechanism for the potential interaction between the microbiome and alloreactivity is not clearly understood, as microbes might influence the host immunity in a multitude of ways. For example, microbial surface antigens can be presented on MHC complexes to be recognized by regulatory T-cells^13,14^. Alternatively, microbial metabolites may affect T-cell differentiation: In particular, metabolites with reported immunomodulatory activity include short-chain fatty acids such as butyrate^15^, nucleosides like inosine^16^, and microbiome-derived secondary bile acids metabolites^17–19^.

In 2023, the standard GVHD prophylaxis strategy for individuals undergoing reduced intensity condition allo-HCT was updated to include a PTCy-based approach^8^. Because most prior studies demonstrating the relationship between the gut microbiome and clinical outcomes were carried out in the pre-PTCy era, we sought to explore whether these associations held in the context of this new prophylaxis approach. Here, we report on the results of the BMT CTN 1801 study, a companion study of BMT CTN 1703. This is the first prospective microbiome-focused study of individuals undergoing allo-HCT within the framework of a multicenter randomized controlled trial.

## Results

### BMT CTN 1801 cohort

In the BMT CTN 1703 study, participants were randomized into two groups: the Tac/MTX arm received the standard GVHD prophylaxis regimen methotrexate and tacrolimus, while the PTCy arm received the experimental prophylaxis consisting of cyclophosphamide at days 3 and 4 post-transplant, followed by mycophenolate mofetil and tacrolimus (see **Fig. 1a**). All participants received reduced intensity conditioning before allo-HCT.

**Figure 1:**
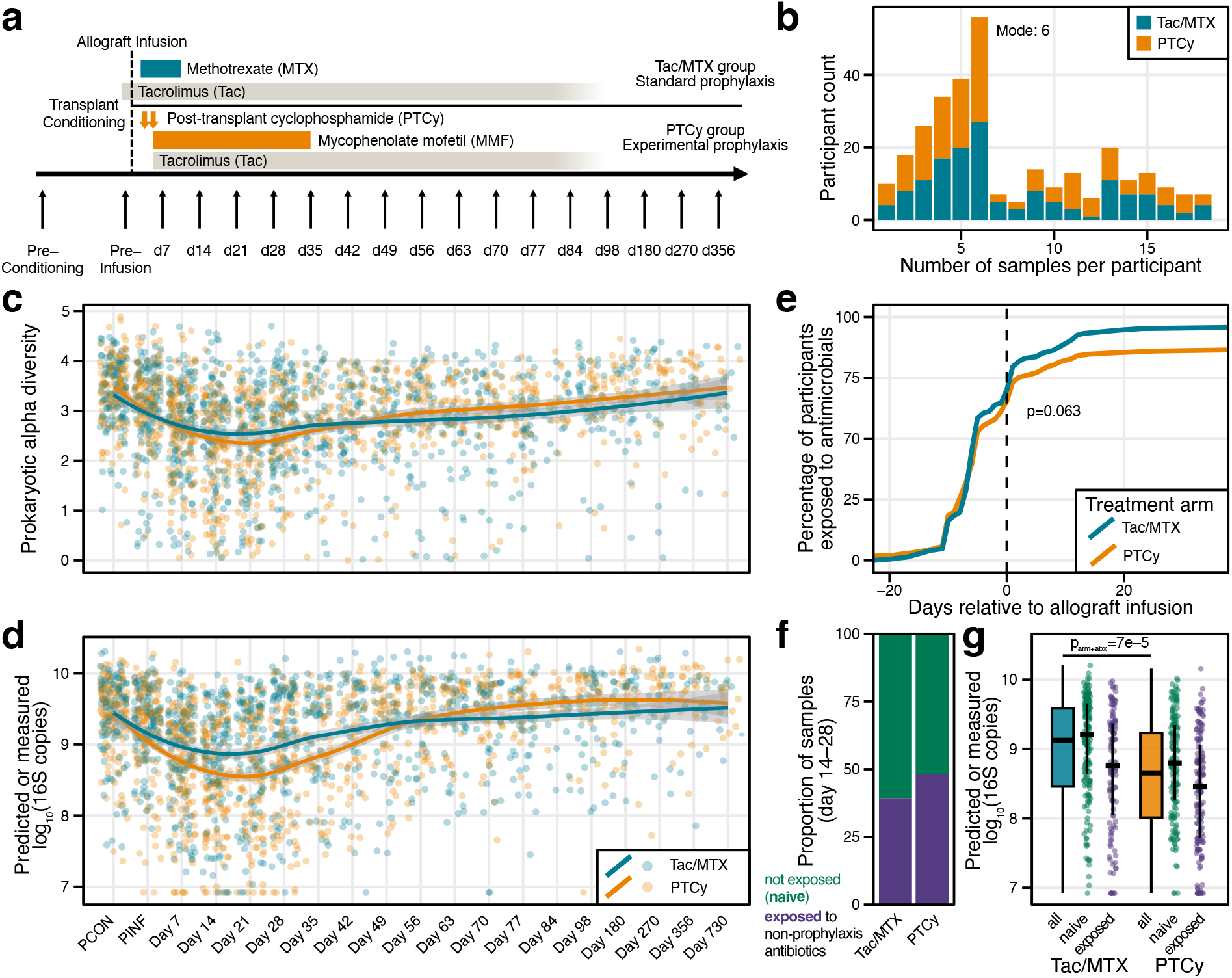
Microbiome diversity and abundance in the BMT CTN 1801 study. **a)** Schematic representation of the BMT CTN 1801 study, showing the GVHD prophylaxis received in the two study arms. Sampling timepoints are indicated by arrows below the timeline. **b)** The number of participants, shown as a bar plot by the number of stool samples they donated over the course of the study. **c)** Microbiome alpha diversity (calculated as Shannon’s index) and **d)** predicted or measured 16S rRNA gene copy number per extraction shown over time for all included stool samples. Samples are binning by collection timepoint and jittered across the x-axis. The trendline for each study arm was estimated using a LOESS model with the formula y∼x, with the grey area showing the 95% confidence interval. PCON: pre-conditioning, PINF: pre-infusion. **e)** Cumulative percentage of participants exposed to any antimicrobial medication up to day 35 post-HCT, split by treatment arm. Difference between distributions was calculated with a log-rank test. **f)** Percentage of stool samples from timepoints 14–28 post-HCT that were either not exposed (naive) or have been exposed to non-prophylaxis antibiotics in the seven days before sampling, shown for both treatment arms as bars. **g)**. Predicted or measured 16S rRNA gene copies per extraction for samples from f), shown as box plots for all samples per treatment arm. Additionally, values are shown as points, colored by non-prophylaxis antibiotics exposure. Thick horizontal bars represent the median within groups. Boxes show the interquartile range (IQR) with vertical lines extending up to 1.5*IQR. For the point clouds, vertical lines represent the IQR. Difference between groups was tested with a linear mixed effect model using GVHD prophylaxis and non-prophylaxis antibiotics exposure as fixed effects and participant ID as random effect.

In total, 324 of the 431 participants enrolled in BMT CTN 1703 co-enrolled in BMT CTN 1801, of whom 304 donated at least one stool sample over the course of the study (Tac/MTX n=147, PTCy n=157, see **SFig. 1**). The most common primary diseases were acute myelogenous leukemia (AML, n=152, 50.0%) and myelodysplastic syndrome (MDS, n=91, 29.9%), with no significant difference between treatment arms (p=0.68, Fisher’s exact test). The only significant difference in other participant characteristics was observed for Donor/Recipient Cytomegalovirus (CMV) status (**Table 1**).

**Table 1:**
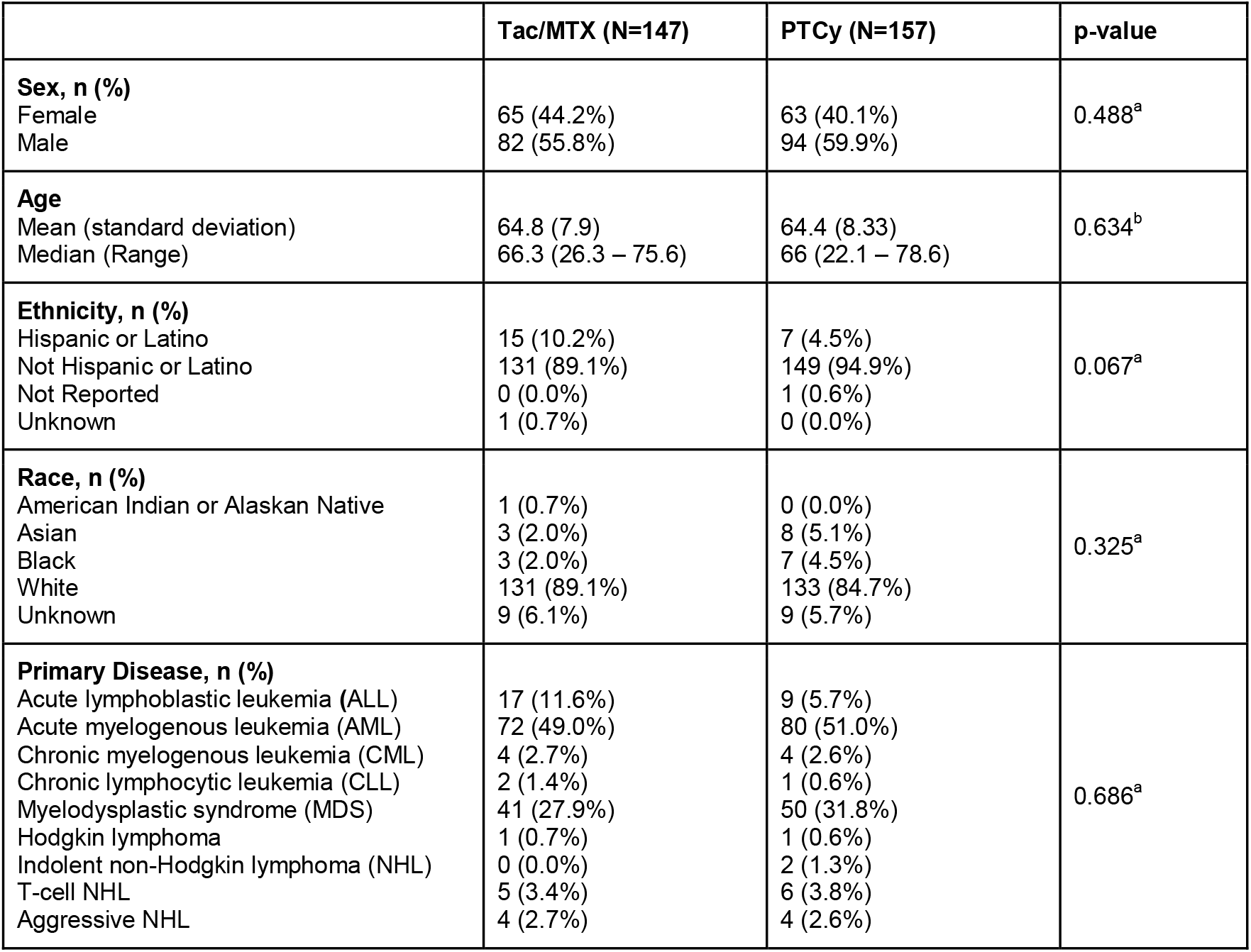
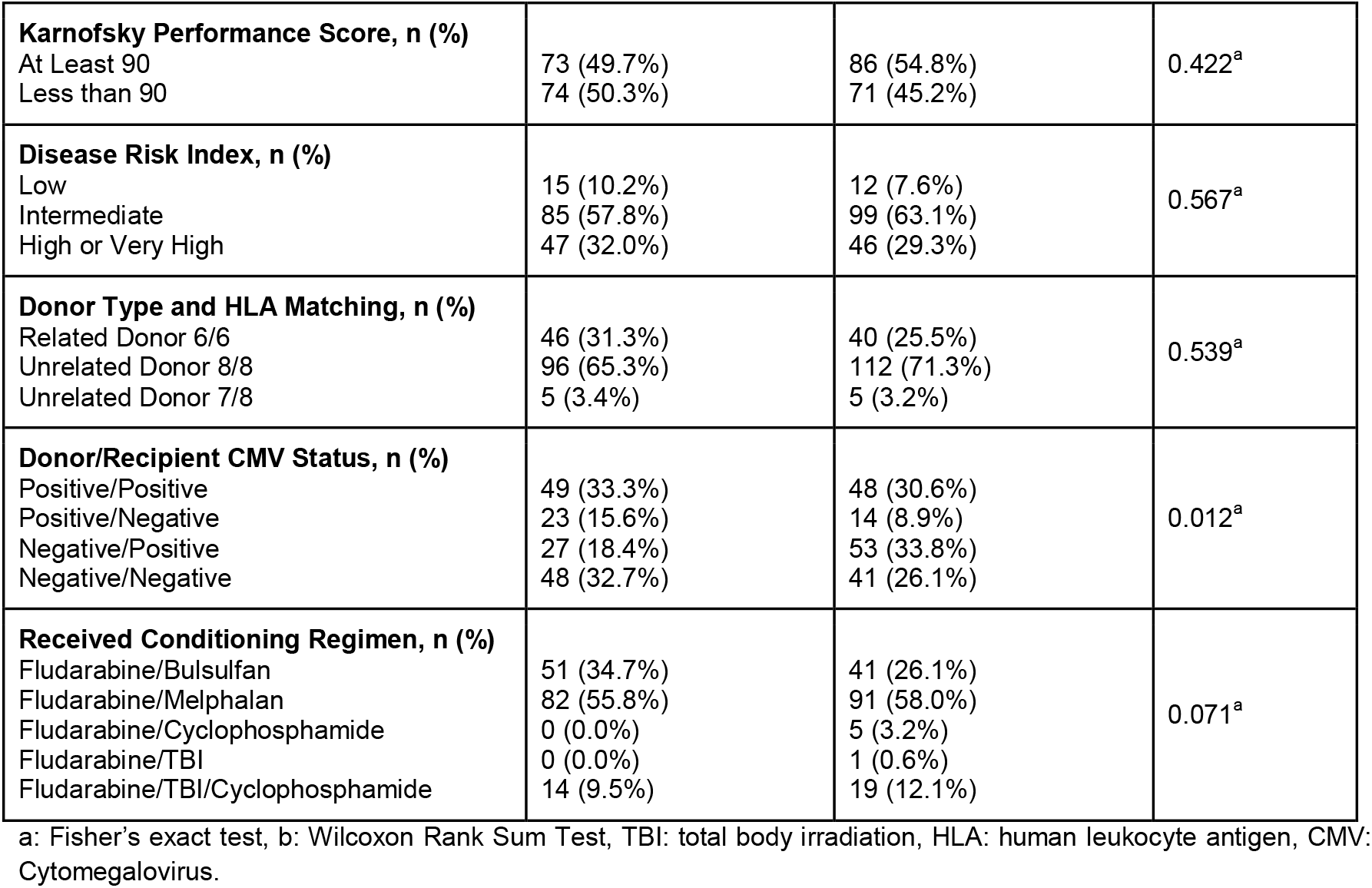
Study population characteristics, overall n = 304.

### Microbiome alpha diversity and absolute microbial load

In total, 2575 stool samples were collected and metagenomically sequenced as part of the trial, with most participants donating 6 samples over the course of their enrollment (**Fig. 1b, SFig. 1**). To first assess the general microbiome state of participants over time, we calculated the microbiome alpha diversity using Shannon’s index and compared the diversity trajectories between treatment arms. At baseline, there was no difference in alpha diversity between groups (p=0.41, Wilcoxon rank sum test). In line with previous reports^11^, diversity declined after allo-HCT in both groups. However, it declined more in the PTCy than in the Tax/MTX arm (**Fig. 1c**). The largest difference between arms was observed between 14 and 28 days post-HCT (p=0.04, linear mixed effect model with participant ID as random effect).

In addition to overall diversity, the absolute microbial load in a sample is hypothesized to be a key ecological factor of importance in the gut microbiome^20^. We therefore determined the number of 16S ribosomal RNA (rRNA) gene copies per DNA extraction, where DNA was extracted from a fixed volume of stool, as a proxy for microbial load. We used a combination of experimental and machine learning techniques^21^ (**Methods, SFig. 2**): the number of 16S rRNA copies were measured directly for 796 samples via digital droplet PCR; the remaining values were predicted using a benchmarked machine learning technique. Similar to diversity, we observed no difference between groups at baseline (p=0.85, Wilcoxon rank sum test), followed by a drop in absolute microbial load after allo-HCT, which was more pronounced in the PTCy group (see **Fig. 1d, SFig. 2**). Between day 14 and 28 post-HCT, the microbial load in the PTCy group was, on average, half that of the Tac/MTX group (p=3e–5, linear mixed effect model with participant ID as random effect).

We initially hypothesized that this observed difference in diversity and absolute microbial load could be driven by differences in antibiotic exposure between groups. In most clinical centers, individuals undergoing allo-HCT are prescribed prophylactic antibiotics at first and – if they develop signs suggestive of infection – are later switched to more broad-spectrum agents. In fact, the initial BMT CTN 1703 study^8^ reported higher incidence of grade 2 infection events in the PTCy group, potentially resulting in the prescription of more antimicrobial agents. However, the number of participants exposed to antibacterials up to day 35 was similar between treatment arms (p=0.063, log-rank test, **Fig. 1e**) and the number of antibacterial doses up to day 35 post-HCT was not significantly higher in the PTCy group (p=0.50, Wilcoxon test, **SFig. 3**). Focusing on individual antibacterial classes, we similarly did not observe significant differences for overall exposure or number of doses that could explain the observed difference in absolute microbial load (see **SFig. 3**).

To further explore the potential impact of antibiotics on absolute microbial load, we compared the number of stool samples collected after participants were exposed to non-prophylaxis antibiotics. We found an excess of non-prophylaxis antibiotic exposure in the PTCy group (p=0.0283, Fisher’s exact test, see **Fig. 1f**), mostly driven by administration of cefepime (see **SFig. 4**). However, adding a term for non-prophylaxis antibiotics to the linear model did not explain the observed difference in microbial load between treatment arms in the 14 to 28 post-HCT timeframe (p=0.0003, linear mixed effect model, see **Fig. 1g**). This suggests that the difference in antibiotic exposure, alone, is not driving the observed difference in absolute microbial load.

Taken together, gut microbiome diversity was lower after allo-HCT, in line with previous reports^11^. We also found that absolute microbial load was decreased in the weeks post-HCT. Both the diversity and absolute load reduction were stronger in the PTCy group compared to the Tac/MTX group, but these divergencies could not be explained by differences in exposure to antimicrobial agents.

### Primary and secondary endpoints: Associations between microbiome diversity and clinical outcomes

The prespecified primary objective of the BMT CTN 1801 study was to evaluate the association between microbiome diversity at neutrophil engraftment and non-relapse mortality (NRM) after one year. The engraftment timepoint was defined as the first of three days that the absolute neutrophil count reached 500 cells/µL blood or higher for 3 consecutive days. Stool samples donated closest after, but within 14 days of the engraftment day were selected for this analysis. On average, engraftment occurred slightly later in the PTCy group (mean=15.3 days) compared to the Tac/MTX group (mean=14.6 days). Microbiome alpha diversity at engraftment was not significantly associated with the cumulative incidence of NRM, either considering diversity categorized by tertiles (p=0.520, Fine-Gray test, see **Fig. 2a**) or treated as a continuous, linear effect (p=0.314, Fine-Gray test, see **Table 2**). Similarly, no significant associations with other clinical outcomes were observed for the microbiome diversity at engraftment (see **Table 2**). GVHD prophylaxis arm was included as a term in the model (see **Methods**), yielding no evidence for differential effects for diversity between treatment arms.

**Table 2:** Association between gut microbiome diversity and clinical outcomes in participants from both arms of the study.

| Clinical outcome | Number of events | Microbiome diversity at engraftment |  | Microbiome diversity at baseline |  |
| --- | --- | --- | --- | --- | --- |
|  |  | P value | Hazard ratio (95% interval) | P value | Hazard ratio (95% interval) |
| **Non-relapse mortality** | **22**<br>[at 12 months] | 0.314<sup>b</sup> | 0.804<br>(0.526, 1.229) | 0.057<sup>b</sup> | 0.631<br>(0.393, 1.014) |
| Relapse/Progression | **62**<br>[at 12 months] | 0.442<sup>b</sup> | 1.092<br>(0.873, 1.365) | **0.047<sup>a</sup>** | 0.389<br>(0.178, 0.849) |
| Acute GVHD [Grade II - IV] | **107**<br>[after 100 days] | 0.060<sup>b</sup> | 0.836<br>(0.693, 1.008) | 0.121<sup>b</sup> | 0.803<br>(0.608, 1.060) |
| Acute GVHD [Grade III - IV] | **18**<br>[after 100 days] | 0.075<sup>b</sup> | 0.710<br>(0.487, 1.035) | **0.032<sup>b</sup>** | 0.610<br>(0.388, 0.958) |
| Chronic GVHD | **60**<br>[at 12 months] | 0.304<sup>b</sup> | 1.130<br>(0.896, 1.425) | 0.119<sup>a</sup> | 2.121<br>(1.018, 4.418) |
| Moderate/severe chronic GVHD | **23**<br>[at 12 months] | 0.333<sup>b</sup> | 1.231<br>(0.808, 1.875) | **0.027<sup>a</sup>** | 3.821<br>(1.233, 11.837) |
| Overall Survival | **55**<br>[at 12 months] | 0.178<sup>b</sup> | 0.846<br>(0.664, 1.079) | **0.007<sup>b</sup>** | 0.631<br>(0.452, 0.881) |
| GVHD-free, relapse-free survival | **124**<br>[at 12 months] | 0.740<sup>b</sup> | 0.972<br>(0.824, 1.148) | **0.047<sup>b</sup>** | 0.786<br>(0.620, 0.996) |

a: Diversity considered as tertiles, b: Diversity considered as linear predictor; model chosen by AIC

**Figure 2:**
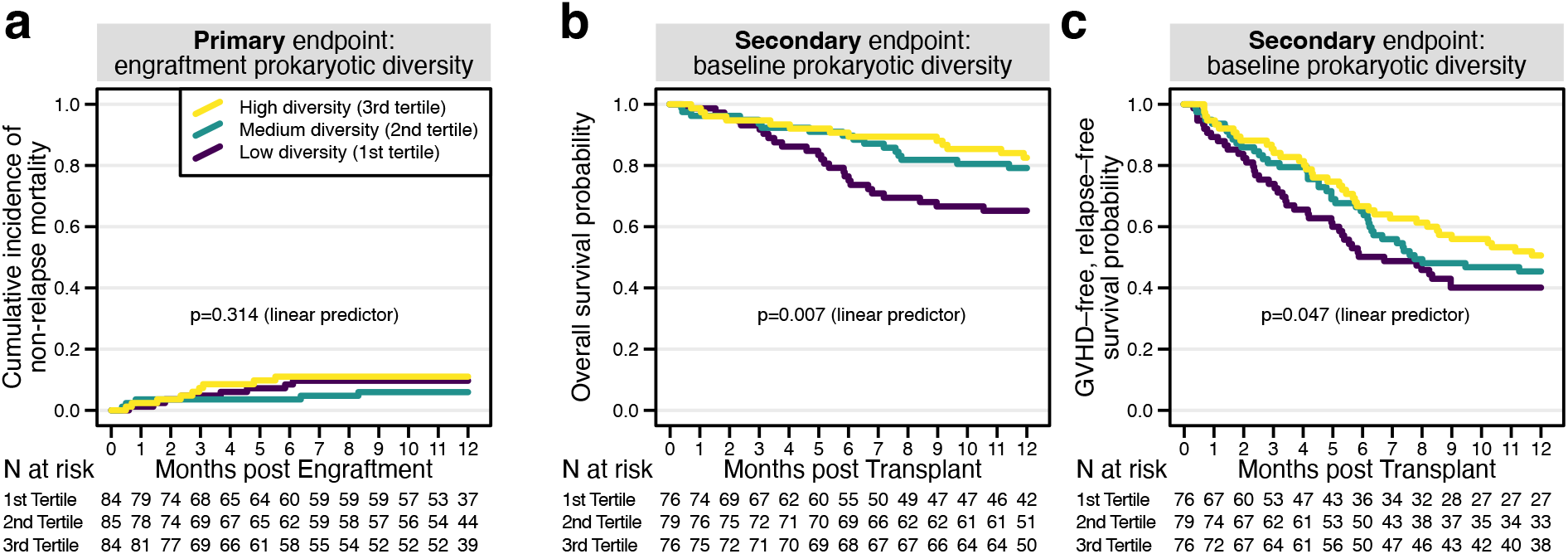
Select association between microbiome diversity and clinical outcomes. **a)** Visualization of the cumulative incidence of non-relapse mortality, separated by the tertiles of microbial diversity at engraftment. **b)** Kaplan-Meier probability curves for overall survival, separated by the tertiles of microbial diversity at baseline. **c)** Kaplan-Meier probability curves for GVHD-free, relapse-free survival (GRFS), separated by the tertiles of microbial diversity at baseline. All p-values were derived from Fine-Gray or Cox regression models.

In a prespecified secondary analysis, we also assessed the association between baseline microbiome diversity (calculated on all pre-conditioning samples) and clinical outcomes. At baseline, we found several clinical outcomes to be associated with microbiome diversity. Compared to the highest diversity tertile, overall survival (OS) was significantly shorter in the lowest tertile diversity group (p=0.031, HR=0.47, Cox proportional hazard model, see **STable 1**). Similarly, lower baseline diversity was associated with shorter OS as a linear predictor (p=0.007, HR=0.63, Cox proportional hazard model, see **Fig. 2b**). The composite primary outcome measure of the BMT CTN 1703 study, GVHD-free, relapse-free survival (GRFS), was similarly associated with baseline microbiome diversity, when treating it as a continuous predictor (p=0.047, HR=0.79, Cox proportional hazard model, **Fig. 2c**).

Another prespecified secondary analysis investigated the association between antimicrobial exposure during the first 30 days post-HCT and clinical outcomes. Participants were dichotomized based on the number of antibacterial doses received (see **Methods**). Similar to microbiome diversity, most outcomes were not strongly associated with antimicrobial exposure.

One notable exception was that the group with higher exposure to antibacterial was at a higher risk for acute GVHD of Grade II-IV (p=0.006, HR=2.6, see **STable 1**), in line with previous reports^22^.

In summary, microbiome diversity at engraftment was not significantly associated with any clinical outcome in this study, in contrast to prior studies^11^. Baseline microbiome diversity, though, was associated with overall survival and other outcomes.

### Microbial domination events

In addition to lower microbial diversity, domination by a single microbial taxon (≥30% of relative abundance explained by a single bacterial genus) is common after allo-HCT^10,11,23^. To quantify microbial domination in the BMT CTN 1801 population, we calculated the rate of domination on species-level over time (see **Fig. 3a**). In agreement with previous studies, microbial domination was commonly observed in both study arms, with ∼50% of samples showing domination at day 14 post-HCT. Despite domination appearing more commonly in the Tac/MTX arm in later timepoints, there was no significant enrichment for dominated samples across GVHD prophylaxis arms over time (p > 0.1 for all time points, Fisher’s exact test).

**Figure 3:**
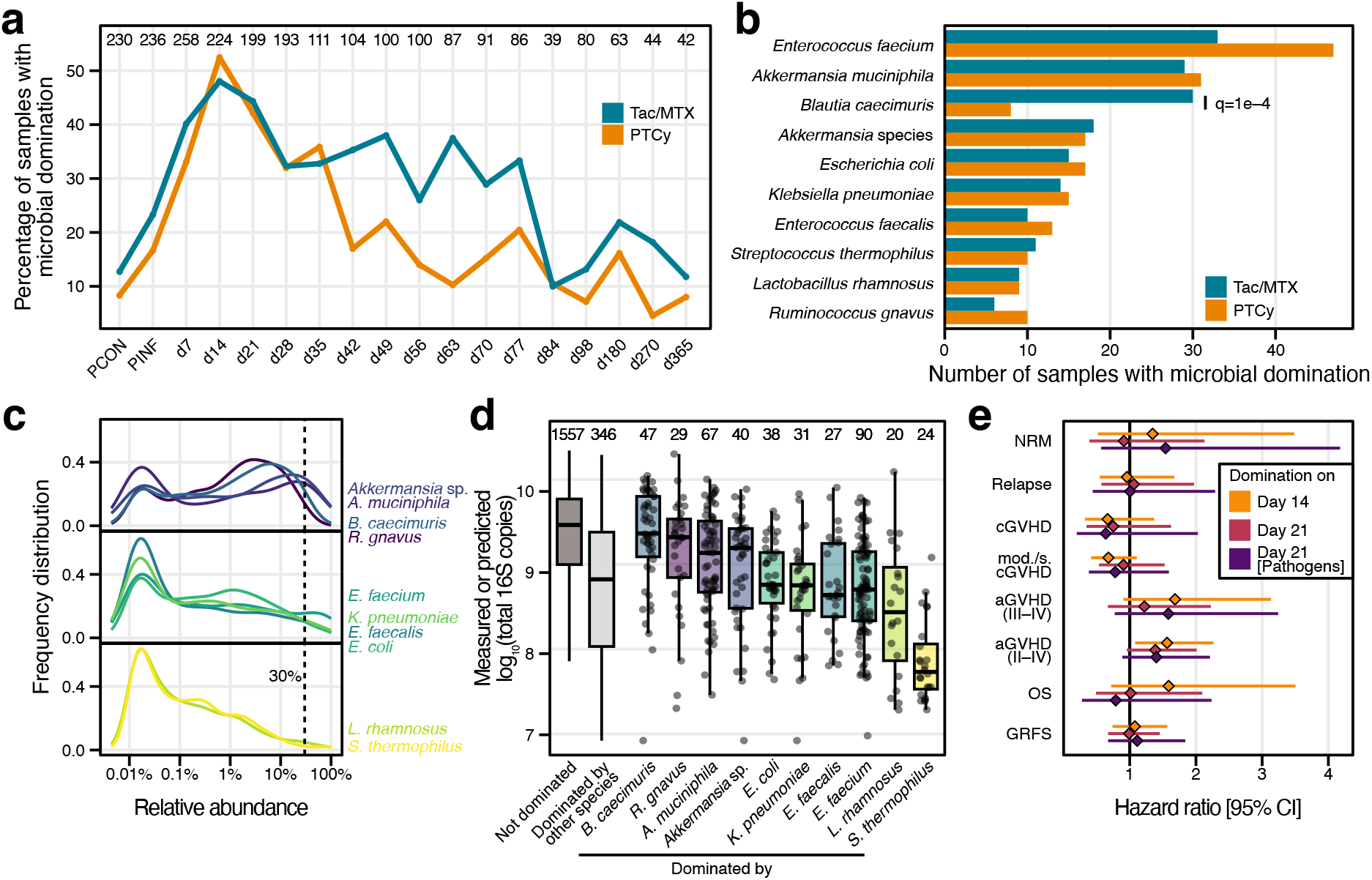
Exploration of microbial domination events. **a)** Percentage of samples showing microbial domination (a single microbial species with relative abundance > 30%) shown for each timepoint and both treatment arms. For all timepoints, differences between treatment arms were not significant as tested by Fisher’s exact test. Numbers on top indicate the number of samples at the given timepoint. **b)** Number of samples with domination by commonly dominating bacteria shown as bars, separated by treatment arm. Differences between treatment arms were tested with Fisher’s exact test; p-values > 0.1 not annotated. **c)** Abundance distribution across all samples for species shown in b). **d)** Predicted or measured 16S copy number per extraction for samples dominated by species shown in b). Numbers on top indicate the number of samples shown. Boxes show the IQR, with vertical lines extending to 1.5*IQR and horizontal thick lines indicating the median. Outliers are not shown. **e)** Hazard ratios (diamonds) with 95% confidence intervals (lines) for Fine-Gray and Cox regression models, investigating the association between microbial domination and clinical outcomes. cGHVD: chronic GVHD, aGVHD: acute GVHD, mod./s.: moderate to severe.

Among the most common bacterial species accounting for domination were pathobionts such as *Enterococcus faecium, Enterococcus faecalis, Escherichia coli*, and *Klebsiella pneumoniae* (see **Fig. 3b**), as well as commensal organisms such as *Ruminococcus gnavus*, a mucin-degrading species observed to dominate the microbiome of individuals with inflammatory bowel disease ^24^. Additionally, two *Akkermansia* species and *Blautia caecimuris* commonly dominated the microbiome, with *B. caecimuris* domination being more common in the Tac/MTX group (q=0.0001, Fisher’s exact test, see **Fig. 3b**). Lastly, *Lactobacillus rhamnosus* and *Streptococcus thermophilus*, two species used in dairy products and probiotics, were also commonly involved in domination events.

To better understand the observed domination events, we investigated the abundance distribution of commonly dominating species (see **Fig. 3c**). Four species (*B. caecimuris, R. gnavus*, and the two *Akkermansia* species) were found to be highly abundant (median relative abundance >1%) in both dominated and non-dominated samples, suggesting that domination by those species might not necessarily be indicative of a highly disrupted microbiome state (see also **SFig. 5**). On the other hand, *L. rhamnosus* and *S. thermophilus* were usually observed at the lower end of the relative abundance spectrum, when present (median relative abundance <0.1%). Compared to samples without domination, samples dominated by those two species also had lower absolute microbial load (see **Fig. 3d**), indicating that domination by those species might not represent microbial blooms, but rather states of community collapse in which microbes provided through the diet become a substantial component of the stool microbiome.

Samples dominated by pathogens fell in the middle of this spectrum: some samples reached both complete domination by a single species as well as comparably high absolute microbial load (see **SFig. 5**), whereas other samples were dominated, yet showed lower microbial load overall. This suggests that some domination events represent microbial blooms by enteric pathogens, while others are likely the consequence of ‘collapse’ of other microbial components of these communities.

Because microbial domination is considered as a ‘dysbiotic’ state that might be indicative of poorer general health, we tested whether microbial domination events were associated with long-term clinical outcomes. We focused on domination at day 14 and 21 post-HCT, as domination was most common on those days, and additionally investigated domination with enteric pathobionts (*E. coli, K. pneumonia, E. faecium*, and *E. faecalis*) at day 21 (see **SFig. 6**). After correction for multiple hypothesis testing, we did not observe significant associations between microbial domination and long-term clinical outcomes (see **Fig. 3e**).

Overall, microbial domination events peaked after allo-HCT, in line with expectations from previous studies, but were not associated with long-term clinical outcomes. Not all domination events involved blooms of enteric pathogens such as *E. faecium*^*10*^, but also microbes with high abundance in non-dominated samples and potentially diet-derived microbes. This suggests that the ecological paths to domination and the likely consequences of domination are varied.

### The gut microbiome differs between treatment arms

To explore the association of GVHD prophylaxis with the microbiome, we identified species with differential abundance between treatment arms (see **Methods**). As expected, no differences were found at baseline (see **SFig. 7**) and similarly, no species were differentially abundant when considering all data across the full study period (see **SFig. 7**).

Focusing on days 14–28 post-HCT, we found several species with lower abundance in the PTCy arm compared to the Tac/MTX arm. The species with the largest difference between treatment arms was *Clostridium scindens* (see **Fig. 4ab**), with other *Clostridia* species showing a similar pattern. The only species with significantly higher abundance in PTCy compared to Tac/MTX, *Streptococcus mutans*, is a predominantly oral-resident bacterium with comparably low relative abundance in stool (see **Fig. 4c**). Other oral taxa also showed a (non-significant) enrichment in PTCy over Tac/MTX (see **STable 3**), potentially reflecting a slight increase in transmission of oral bacteria to the gut^25^ in the PTCy arm. The difference in several Clostridial species, including *C. scindens*, was not explained by non-prophylaxis antibiotics (see **SFig. 7**), suggesting an effect of GVHD prophylaxis on the microbiome that might be specific to organisms of this clade.

**Figure 4:**
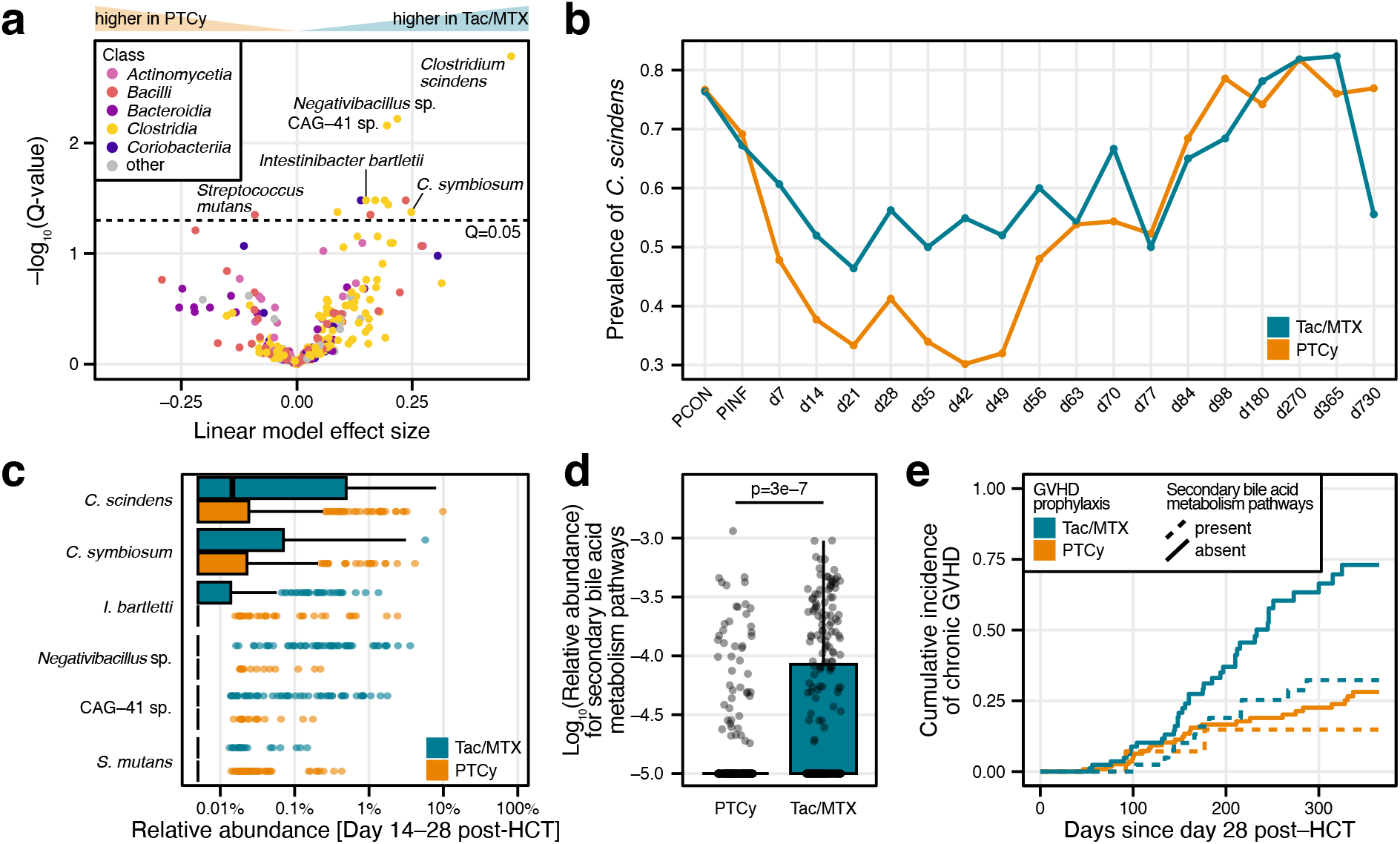
Differences in microbiome composition and functional capacity linked to GVHD prophylaxis. **a)** Volcano plot showing differential abundance between PTCy and Tac/MTX in the days 14–28 post-HCT. Linear mixed effect models were constructed with Participant ID as random effect. Dots represent individual microbial species, colored by their GTDB class annotation. **b)** Prevalence trajectory for *C. scindens* across study arms. **c)** Log-transformed relative abundance distribution for species highlighted in a), separated by study arm. Boxes show the IQR, with vertical lines extending to 1.5*IQR and horizontal thick lines indicating the median. Outliers are shown as dots. **d)** Log-transformed relative abundance of secondary bile acid metabolism pathways on days 14–28 post-HCT, separated by study arm. Boxplot definition as in c); all data are shown as dots. **e)** Visualization of the cumulative incidence for chronic GVHD, separated by study arm and presence or absence of secondary bile acid metabolism pathways at days 14–28 post-HCT. Association between chronic GVHD and group (study arm and secondary bile acid presence) was tested using a Fine-Gray model with death as competing risk.

*C. scindens* is known to have the capacity to modify host-derived primary bile acids into secondary bile acids via the *bai* operon^26^. In particular, *C. scindens* can perform 7α/β-dehydroxylation of primary bile acids, resulting in deoxycholic and lithocholic acid. Secondary bile acids and their myriad derivatives are regulators of host immune states, acting both on the innate and adaptive immune systems by inducing the differentiation of regulatory T cells^27^.

We therefore asked if the lower prevalence of *C. scindens* was also reflected in a functional, gene-based analysis of the metagenomic sequencing data (see **Methods**). Interestingly, secondary bile acid metabolism pathways were the only pathways with significantly lower abundance in the PTCy arm at days 14–28 post-HCT at q < 0.01 (see **Fig. 4d** and **SFig. 7**). There is incomplete correspondence between the presence of secondary bile acid metabolism pathways and of *C. scindens*: the detection of metabolic pathways appears to be less sensitive, but more specific, than the quantification of *C. scindens* abundance from taxonomic profiling (see **SFig. 8**).

Given the ability of bile acids to modulate the host’s immune system, we asked if the presence of *C. scindens* or of secondary bile acid metabolism pathways at days 14–28 post-HCT was associated with clinical outcomes. The presence of *C. scindens* was not associated with clinical outcomes (see **STable 4**), whereas the presence of secondary bile acid metabolism pathways during day 14–28 was associated with lower risk for developing chronic GVHD (see **Fig. 4e, SFig. 8**, and **STable 4**). Interestingly, the presence of secondary bile acid metabolism pathways corresponded to lower chronic GVHD risk mostly in the Tac/MTX arm, resulting in chronic GVHD incidence similar to PTCy-based GVHD prophylaxis.

This analysis revealed that PTCy and Tac/MTX treatment are associated with different states of microbiome disruption at days 14–28 post-HCT, with participants in the PTCy arm exhibiting lower prevalence of potentially beneficial *Clostridia* species, especially bile-acid modifying *C. scindens*. The association between secondary bile acid metabolism pathways and chronic GVHD risk was observed predominantly in the Tac/MTX group. This suggests that a potential modulation of T-cell alloreactivity by microbiome-derived secondary bile acids might be less relevant for chronic GVHD prevention in conjunction with PTCy treatment.

## Discussion

The BMT CTN 1801 study is one of the largest, prospective randomized controlled clinical trials to collect stool samples for microbiome analyses. In line with previous work, we observed microbiome disruption after allo-HCT, which intriguingly differed across the two GVHD prophylaxis arms. Associations between microbial diversity and clinical outcomes were observed for baseline, but not for engraftment microbiome. Treatment arms exhibited a similar number of total microbial domination events, yet showed significant differences in *Clostridia* species related to secondary bile acid metabolism.

In comparison to Tac/MTX, PTCy treatment was associated with lower alpha diversity, lower absolute microbial load, and lower prevalence of *Clostridia* species at days 14–28 post-HCT, none of which could be explained by differential antibiotic exposures. Since PTCy was administered on days 3 and 4 post-HCT and since cyclophosphamide and its metabolites are usually cleared through the urine within 24 hours after infusion^28^, a direct effect of PTCy on the gut microbiome seems unlikely. In mice, cyclophosphamide treatment is similarly associated with a delayed effect on the gut microbiome composition (one week after treatment), yet resulted in immediate alterations in the gut epithelial barrier^29^. Therefore, PTCy’s impact on the microbiome might be explained by an effect on gut epithelial cell homeostasis and the integrity and cellular composition of the colonic lamina propria^30^, even though we did not observe an enrichment of human reads in the stool samples of the PTCy group (see **SFig. 2**).

A confounding factor for interpretation of these results is that MMF was additionally administered in the PTCy arm up to day 35 post-HCT. MMF is a prodrug for mycophenolic acid, which undergoes enterohepatic circulation and is associated with gastrointestinal toxicity as a common side effect^31^. Therefore, some of the observed differences could be due to direct effects of MMF on the gut epithelium or on the gut microbiome. On the other hand, differences between the Tac/MTX and PTCy arms were less pronounced at day 35 (see **Fig. 1**) and absolute microbial load was already trending upwards on day 28 (see **Fig. 1**), arguing against a strong direct effect of MMF.

In our prespecified analyses, microbiome diversity at engraftment was not associated with non-relapse mortality and other clinical outcomes (see **Table 2** and **Fig. 2**). These findings challenge the established link between low gut microbiome diversity and mortality in allo-HCT^11^, especially given that PTCy was associated with lower diversity than Tac/MTX, yet better clinical outcomes^8^. Differences in reported associations might be explained by technical differences between studies, such as sequencing strategy (16S rRNA amplicon versus shotgun metagenomic sequencing), analysis framework (choice of diversity index, taxonomic resolution, and timepoint for analysis), or clinical parameters (reduced intensity compared to myeloablative conditioning). However, the study by Peled et al.^11^ reported that the associations between clinical outcomes and microbiome diversity were not seen in T-cell-depleted grafts. The mechanism by which PTCy treatment is believed to prevent GVHD is similar to T-cell depletion, in that dividing T-cells, including allo-reactive T-cells, are inactivated by PTCy, whereas non-allo-reactive graft T-cells and regulatory T-cells are spared to provide immune reconstitution in the long term^7^. This suggests that microbial antigens or metabolites might contribute to the regulation of allo-reactive T-cells, a cell population that is efficiently ablated by PTCy.

Supporting this model is the finding that *C. scindens* and bile-acid metabolism pathways were found to be of lower prevalence in the PTCy arm compared to the Tac/MTX arm. Secondary bile acids are implicated in the differentiation of regulatory T cells and Th17 cells ^17–19^, which in turn can regulate allo-reactive T-cells. Similarly, the pool of fecal bile acids are altered in participants with auto-immune diseases, including inflammatory bowel disease^32^, or in participants undergoing allo-HCT^33^. Here, lack of secondary bile acid metabolism pathways was associated with an increased risk for chronic GVHD in the Tac/MTX group, but not in PTCy, highlighting again that microbial contributions might be less relevant in conjunction with PTCy-based GVHD prevention.

While our analyses did not fully recapitulate the established associations between microbiome diversity and clinical outcomes for the engraftment timepoint, they raise several interesting areas for follow-up studies. First, we observed a strong association between baseline microbiome diversity and overall survival. The baseline microbiome composition of allo-HCT recipients might reflect additional risk factors such as prior antibiotic or chemotherapy use that should be explored further. Secondly, the presence of bile acid metabolism pathways might be important for the development of chronic GVHD (see **STable 4**), proposing mechanistic follow-up studies. Lastly, in contrast to prior studies that mostly focused on domination of the gut microbiome by pathogens such as *E. coli* or *Enterococcus* species^10,23^, we find several instances of domination with non-pathogen species in individuals receiving reduced intensity conditioning. Thus, a more nuanced understanding of the relationship between microbiome domination and clinical outcomes is likely warranted.

This study has several limitations. First, the stool sample chosen for the engraftment timepoint might not accurately represent the engraftment microbiome, as a maximum of 14 days between neutrophil engraftment and sample collection were allowed. In 11% of participants, the stool sample had been collected 10 or more days after neutrophil engraftment. Additionally, for 34 participants, the absolute neutrophil count never declined below 500 cells/µL. Following the prespecified analysis plan, the engraftment timepoint was set to the first day post-HCT, possibly resulting in the selection of an inappropriate, early stool sample. Second, part of the absolute microbial load data was predicted with a machine learning model trained on the observed correlation between total DNA concentration and 16S copy number (see **SFig. 2**, ref.^21^). While we generated additional measurements, samples with low abundance or domination by fungal pathogens^34^ might be challenging to predict accurately. Third, the bile acid pool in the stool of participants was not directly measured in this study. Instead, we measured the genes encoding the enzymes of these metabolic pathways in the microbiome and inferred an altered bile acid pool. Future studies will be needed to elucidate the role of fecal bile acid concentration and composition for GVHD risk.

## Conclusion

This study provides intriguing glimpses into the complex interplay between the gut microbiome, GVHD prophylaxis, and the immune system in participants undergoing allo-HCT and challenges some prevailing models of the role for the microbiome in clinical allo-HCT outcomes. While PTCy was associated with better one-year clinical outcomes in BMT CTN 1703^8^, the long-term consequences of the PTCy-associated strong suppression of the microbiome should be investigated with longer follow-up. Additionally, mechanistic studies are needed to disentangle the direct and indirect interactions between T-cell repertoire, PTCy, and the microbiome.

## Methods

### Clinical trial design

The BMT CTN 1801 clinical trial was a companion study to the BMT CTN 1703 trial^8^, a Phase 3 multi-center randomized controlled clinical trial to compare the efficacy of two GVHD prophylaxis regimens, the standard prophylaxis (tacrolimus and methotrexate) and an experimental prophylaxis (post-transplant cyclophosphamide, tacrolimus, and mycophenolate mofetil). Information about the clinical trial is available under https://clinicaltrials.gov/study/NCT03959241.

The primary objective of the BMT CTN 1801 study was to evaluate the association between the microbial diversity at engraftment with non-relapse mortality at one year. Participants were co-enrolled into BMT CTN 1801 with a goal of 300 participants (70% of the 428 participant enrollment goal for BMT CTN 1703). No major differences in terms of demographic or other characteristics were observed between the full BMT CTN 1703 study population and the BMT CTN 1801 subset populations (see the companion manuscript describing T-cell receptor sequencing, ref.^35^). Informed consent was obtained from all participants. The study protocol was approved by the Institutional Review Board of the National Marrow Donor Program (NMDP). This manuscript was submitted on behalf of the Blood and Marrow Transplant Clinical Trials Network (BMT CTN).

### Sample collection and processing

Stool samples were collected before conditioning (PCON), before infusion of hematopoietic cells (PINF), and at weekly intervals after infusion up to day 84 post-HCT, followed by less frequent sampling (day 98, 180, 270, 365 and 730 post-HCT). All samples were collected without preservatives. Most samples were stored at 4°C within 30 min after collection and then shipped overnight to the NMDP, where they were frozen at −80°C. A subset of samples (n=250) were frozen directly after collection. No major differences in composition or diversity were observed between same-day versus next-day sample storage (see **SFig. 9**).

In total, 2575 stool samples were collected. From every frozen sample, we aimed to extract a consistent amount of stool (∼200mg) with a biopsy punch. Then, DNA was extracted using the QIAamp PowerFecal Pro DNA Kit according to the manufacturer’s instructions and eluted into 100µL elution buffer onto 96-well plates.

The origin of one sample tube could not be verified, resulting in the removal of the resulting sequencing data (see **SFig. 1**).

### Metagenomic shotgun sequencing

For sequencing, 30µL of extracted DNA were used as input for metagenomic library preparation with the NEB Ultra II kit, according to the manufacturer’s instructions. DNA concentration and DNA quality was measured using an Agilent 5400 Fragment Analyzer System prior to sequencing. Each 96-well plate of DNA extractions contained a positive (ZymoBIOMICs microbial community standard) and a negative extraction control (water). Finally, libraries were pooled and sequenced with the NovaSeq 6000 platform, generating 2 × 150 bp reads with a target depth of 6 Giga-basepairs total sequencing (see **STable 2**).

### Metagenomic data processing

Raw reads were processed with a standard workflow implemented in NextFlow v23.04.3^36^, available under github.com/bhattlab/bhattlab_workflows_nf. In short, raw paired-end reads were deduplicated with HTStream SuperDeduper v1.3.3 before removing low-quality bases and short reads with TrimGalore v0.6.7. Quality-controlled reads were mapped against the human genome (version hg38) using bwa v0.7.17^37^ and all matching reads were discarded.

Taxonomic profiling was performed with mOTUs v3.0.3 ^38^, using the parameters -g 2 -l 75. Samples with low sequencing quality (less than one million reads after host removal and a count from the motus profiler less than hundred) were removed from the analysis (see **SFig. 1**).

Microbiome alpha diversity was calculated as Shannon’s index, using the vegan v2.7-1 R package on samples rarefied to 3000 motus counts.

Functional profiles were generated with HUMAnN v3.9 (ref ^39^), using default parameters.

### Quantification of absolute microbial load

To quantify the absolute microbial load in each sample, we performed digital droplet PCR (ddPCR) for the 16S rRNA gene, as previously described^21,40^. After running the experiments on a subset (∼20%) of samples, we observed a strong correlation between DNA concentration and 16S copy number per DNA extraction, prompting us to develop a machine learning model for absolute microbial load prediction (see ref.^21^).

The initial subset of samples measured via ddPCR skewed towards higher DNA concentration compared to the overall distribution of DNA concentration observed in our samples (see **SFig. 2**). Therefore, we additionally measured the 16S copy number via ddPCR in four additional 96-well plates to better represent the spread of DNA concentration in the measured subset. The ddPCR experiments were performed as described in ref.^40^ as well.

Given the high accuracy of the model (concordance correlation coefficient of 0.91), the 16S copy number per extraction was estimated for the remaining samples with the trained machine learning mode (see **SFig. 2**).

### Statistical analysis of microbiome diversity and absolute microbial load

To test for differences in microbiome diversity and absolute microbial load between GVHD prophylaxis arms, we used a Wilcoxon test for single timepoints. When multiple timepoints were considered, a linear mixed effect model was used with participant ID as random effect. P-values were calculated with the lmerTest package^41^.

### Quantifying exposure to and the effects of exposure to antibiotics

To quantify differences in antimicrobial exposure between GVHD prophylaxis arms up to day 35, we determined the first date of antibiotic exposure for each participant. The difference between GVHD prophylaxis arms in initial exposure was tested with a log-rank test through the survival package in R. If a participant was exposed to antibacterials only after day 35, they were censored for this analysis. Additionally, the number of prescribed antibiotics was summarized for each participant and differences between arms were tested using a Wilcoxon test. Similarly, this analysis was repeated for each class of antibiotics.

To explore the effect of prophylaxis and non-prophylaxis antibiotics on the microbiome up to day 35 post-HCT, we asked for each stool sample whether the participant had been exposed to antibiotics within seven days before donating the sample. We focused on antibiotics for which at least ten samples were considered to be exposed and constructed linear models to estimate the effect of drug treatment on the 16S copy number per extraction. Each antibiotic (together with the route of application) was classified as predominantly prophylactic or non-prophylactic therapeutic, based on common protocols. This approach might mis-classify some antibiotic applications, as treatment plans differ between participating hospitals, but should capture broad trends of antibiotic application.

### Statistical analysis for the primary and secondary study objective

The statistical analysis was performed according to the statistical analysis plan. The primary object was to evaluate the association between microbiome diversity at neutrophil engraftment and non-relapse mortality (NRM) after one year. The target sample size for the study was determined to provide at least 85% power to detect a 20% difference in the cumulative incidence of NRM between any two microbiome diversity tertile groups using pairwise comparisons. The secondary objectives encompassed the evaluation of associations between i) microbiome diversity at neutrophil engraftment and other clinical outcomes, ii) microbiome diversity at baseline and clinical outcomes, and iii) between antimicrobial exposure during the first 30 days post-HCT and clinical outcomes.

To evaluate the associations between microbiome alpha diversity and clinical outcomes, diversity at the time of neutrophil engraftment was classified into tertiles. For each participant, the stool sample directly at or within 14 days of engraftment was identified for this analysis. For the analysis of the baseline diversity, samples before conditioning (PCON) were chosen.

To compare the effects of diversity groups on the cumulative incidences of NRM, grade II-IV and III-IV acute GVHD, all grade and moderate/severe chronic GVHD, and relapse/progression, Fine-Gray regression models were used with adjustment for the GVHD prophylaxis group. An additional set of Fine-Gray models included microbiome diversity as a continuous variable in a Fine-Gray regression model while adjusting for GVHD prophylaxis. A linear diversity effect was used since investigation of the martingale-type residuals from the Fine-Gray models indicated either no clear trend or that a linear effect was appropriate. Additional baseline variables including age, gender, race/ethnicity, Karnofsky performance status (<90 vs. >=90) at transplant, primary disease at transplant, disease risk index (DRI), hematopoietic cell transplant comorbidity index (HCT-CI), donor type and HLA matching, donor/recipient CMV status, donor/recipient sex match, donor/recipient ABO match, and conditioning regimen were considered for inclusion into the models using stepwise selection with a p-value of less than 0.05 as the selection criterion. Due to the small number of events for non-relapse mortality (n=22), acute GVHD of grade III-IV (n=18), and moderate/chronic GVHD (n=23), no additional variable selection was considered in these models.

For overall survival (OS) and GVHD-free, relapse-free survival (GRFS), Cox regression models were constructed similarly. Events for OS were defined as death from any cause; for GRFS, events included death from any cause, acute GVHD grade III-IV, chronic GVHD onset requiring immunosuppression, and relapse or progression of the primary disease.

Finally, the Akaike information criterion (AIC) was used to assess the predictive ability of the Fine-Gray and Cox regression models. To select between models considering diversity either as tertiles or as linear predictor, the model with the lower AIC was chosen. All analyses were performed in SAS version 9.4 and results were visualized in R version 4.0 or higher.

To quantify the association between exposure to antimicrobials and clinical outcomes, the number of antimicrobial medication dose-days up to day 30 post-HCT was determined for each participant. A participant who received two different antimicrobials for one day was recorded as having received two dose-days, whereas a participant who received one antimicrobial for three days was recorded as having received three dose-days. For combination drugs, dose-days were determined individually for each of the two components of the combination therapy. Participants were dichotomized into high or low antimicrobial exposure based on the median number of dose-days across all participants. Fine-Gray and Cox regression models were constructed as described above.

### Domination events

For the exploration of microbial domination events, we defined samples to be dominated, if a single microbial species was present at more than 30% relative abundance. Differences in domination rate between treatment arms were tested using Fisher’s exact test. In-depth analysis was restricted to microbial species most commonly resulting in domination, with more than 20 samples dominated by the species in question. Association between domination on day 14, on day 21, or domination by potential pathogen (*E. coli, K. pneumoniae, E. faecium*, and *E. faecalis*) on day 21 post-HCT (maximum rate of domination by pathogens, see **SFig. 6**) were tested with Fine-Gray or Cox regression models as described above.

### Differential abundance testing

To test for differential abundance of microbial species between treatment arms, we tested log-transformed relative abundance values with a mixed linear effect model using GVHD prophylaxis as fixed effect and participant ID as random effect. P-values were calculated using the lmerTest package^41^. For testing baseline differences, a standard linear model without random effects was used, as no repeated samples for the same participant were included. To account for multiple hypothesis testing, p-values were corrected using the Benjamini-Hochberg procedure^42^.

## Acknowledgements

We acknowledge all members and participating clinical centers of the BMT CTN 1703/1801 trial and all participants that generously donated samples for the study. We further thank the members of the Bhatt lab, especially M.P. Grieshop, for valuable feedback.

This manuscript was prepared using BMT CTN 1703/1801 Research Materials obtained from the BMT CTN Repository operated by the NMDP. Support for this study was provided by grants #U10HL069294 and #U24HL138660 to the BMT CTN from the National Heart, Lung, and Blood Institute (NHLBI) and the National Cancer Institute (NCI). The Center for International Blood and Marrow Transplant Research (CIBMTR) is supported primarily by U24-CA076518 from the NCI, the NHLBI, and the National Institute of Allergy and Infectious Diseases and by contract HHSH234200637015C to the CIBMTR from HRSA/DHHS. The content is solely the responsibility of the authors and does not necessarily reflect the opinions or views of the BMT CTN 1703/1801 protocol team, the BMT CTN, CIBMTR, the NHLBI, or NCI. We acknowledge support from the Biostatistics Shared Resource at the Medical College of Wisconsin Cancer Center.

J.W. is a Damon Runyon Quantitative Biology Fellow supported by the Damon Runyon Cancer Research Foundation (DRQ-22-24). T.M.A is supported by NIH K23 AI163365. T.F. acknowledges funding from the P30-CA008748 MSK Cancer Center Support Grant/Core Grant. J.U.P, M.A.P. and T.F. are supported in part by the NIH/NCI Cancer Center Support Grant P30 CA008748. B.D. was supported by the Stanford Medical Scholars Fellowship Program, Stanford Berg Scholars Program and a Physician Scientist Institutional Award (PSIA) from the Burroughs Wellcome Fund. N.B.P. is supported by NIH T32 5T32AI007502. A.D.S. and collection for some of the samples for this study is supported by NIH R01AG066719 and R01HL151365. L.S.K. is supported by National Institutes of Health UG1HL174426, U19AI174967, P01 HL158505, R01HL095791, P01HL158504, and R01HL181969. A.S.B. is supported by the Allen Distinguished Investigator award and the Bhatt lab is supported by NIH R01AI148623, R01AI143757, R01HL181969, and R01CA301727, and a Stand Up 2 Cancer Grant.

## Author contributions

E.F.B., B.D., J.W. – Sample handling and processing, ddPCR experiments

J.W., M.J.M., T.M.A., J.U.P., N.B.P., T.F., W.S., L.S.K., M.A.P., A.S.B. – Data analysis and interpretation

H.E., M.G., M.M.A.M., A.R.R., L.R. – PIs of top accruing centers

W.S. – Study oversight, BMT CTN 1801 protocol officer

J.U.P., T.M.A., R.R.J, M.B., B.B., S.C., S.W.C., W.C., S.D., H.E., M.G., R.J.J, J.E.L., B.R.L., M.M.A.M., H.S.M., A.R., A.R.R., M.L.R, L.R., K.S., A.S., A.D.S. – Study design, BMT CTN 1801 protocol members

M.M.H., M.H., S.G.H., J.B.M. - BMT CTN leadership / companion protocol 1703 protocol chairs

L.S.K., M.A.P., A.S.B – Study design, BMT CTN 1801 protocol chairs

M.R.M.vdB., L.S.K., M.A.P., A.S.B. – Conceptualization

J.W., A.S.B., J.U.P. - writing the original manuscript

All authors - editing and approving the final manuscript

## Competing interests

J.U.P reports research funding, intellectual property fees, and travel reimbursement from Seres Therapeutics, and consulting fees from DaVolterra, CSL Behring, Crestone Inc, MaaT Pharma, and Canaccord Genuity, Inc and RA Capital. He serves on an Advisory board of and holds equity in Postbiotics Plus Research. He serves on an Advisory board of and holds equity in Prodigy Biosciences. He has filed intellectual property applications related to the microbiome (reference numbers #62/843,849, #62/977,908, and #15/756,845). Memorial Sloan Kettering Cancer Center (MSK) has financial interests relative to Seres Therapeutics.

T.M.A. is a paid consultant for Seres Therapeutics LLC.

M.H. reports research support from ADC Therapeutics, Spectrum Pharmaceuticals, and Astellas Pharma. He received consulting fees from Caribou, Autolus, Forte Biosciences, Byondis, Daiichi Sankyo, BMS, Caribou, Kite, Incyte, and Abbvie, serves on the Speaker’s Bureau for AstraZeneca, ADC Therapeutics, BeiGene, Kite, Sobi, and on the DMC for Myeloid Therapeutics (2023), CRISPR (2024).

A.D.S. has received research funding from Merck, Novartis, Enterome, and Seres; research product from DSM/iHealth, Clasado, and BlueSpark Technologies; and consulted for Targazyme, Acrotech, Geron, Janssen, Beigene, Seres, BVF, and GSK.

H.S.M. reports consulting activity and serving on scientific advisory boards for Geron, Ascentage, Abbvie, Syndax Therapeutics, Kite Gilead, Seres, Sobi, BMS, Incyte, and Sanofi.

S.C. received honoraria from Sanofi, Sobi, Ascentage Pharma, BMS, Pfizer, Legend Biotech; Institutional Research Funding: C4 Therapeutics, CARSgen, Cynata Therapeutics, Johnson & Johnson, AstraZeneca, Incyte, Abbvie, and Cullinan Therapeutics.

S.G.H. discloses competing interests for VITRAC Therapeutics (research funding), Incyte (research funding), CSL Behring (clinical trial adjudication), Sanofi (non-branded educational programming), MaaT Pharma (advisory board), and Ossium (advisory board)

A.R.R. served as medical expert witness for the US Department of Justice.

L.S.K. is a Scientific founder and has equity in Regatta Bio. She is a member of the Scientific advisory board for HiFiBio. She has research funding from Tessera Therapeutics, EMD Serono, Gilead Pharmaceuticals, and Regeneron Pharmaceuticals. She receives consulting fees from Vertex and Bluerock Therapeutics. She receives grants and personal fees from Bristol Myers Squibb. Her conflict-of-interest with Bristol Myers Squibb is managed under an agreement with Harvard Medical School.

M.A.P. reports honoraria from Allogene, Celgene, Bristol-Myers Squibb, Exevir, ImmPACT Bio, Incyte, Kite/Gilead, Merck, Miltenyi Biotec, Nektar Therapeutics, Novartis, Omeros, OrcaBio, Pierre Fabre, Sanofi, Syncopation, Takeda, VectivBio AG, and Vor Biopharma. He serves on DSMBs for Cidara Therapeutics and Sellas Life Sciences. He has ownership interests in Omeros and OrcaBio. He has received institutional research support for clinical trials from Allogene, Genmab, Incyte, Kite/Gilead, Miltenyi Biotec, Novartis, and Tr1x.

A.S.B. is a Founder of Stylus Medicine and serves on the scientific advisory board and is a board observer. She also serves on the Scientific Advisory Board of Caribou Biosciences and Cantata Biosciences.

## Data availability

The raw sequencing data for all samples sequenced in this study are available from the European Nucleotide Archive under the study identifier <u>PRJEBXXXX</u>.

## Code availability

Source code for analysis and figure generation is publicly available at Zenodo under https://doi.org/10.5281/zenodo.17127362 and on GitHub under https://github.com/jakob-wirbel/BMT_CTN_1801.

