## Supplemental Figures for "Differential effects of two common GVHD prophylaxis regimens on the gut microbiome: Results from the BMT CTN 1801 study"

**Supplementary Figures**

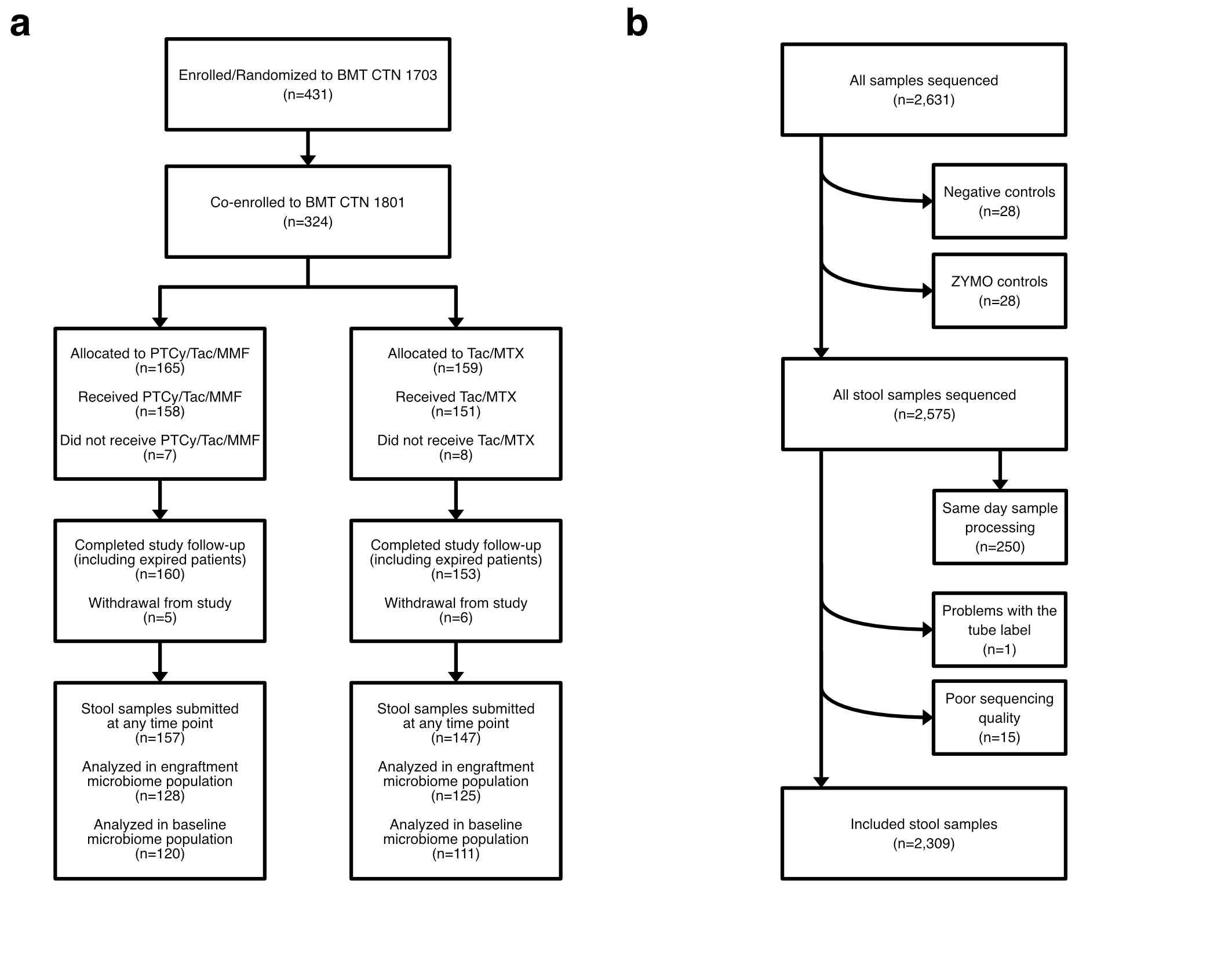

**SFig. 1: Flowcharts accounting for participant enrollment and stool sample processing**

**a)** Flowchart detailing the number of participants that co-enrolled in BMT CTN 1801, provided stool samples, and were included in the primary and secondary pre-specified analyses. **b)** Overview over all sequenced samples and the filtering steps employed to arrive at the final set of sequenced stool samples. Samples that underwent same-day processing (see Methods) were excluded based on limited differences compared to matched samples that were processed on the next day (see **SFig. 9**).

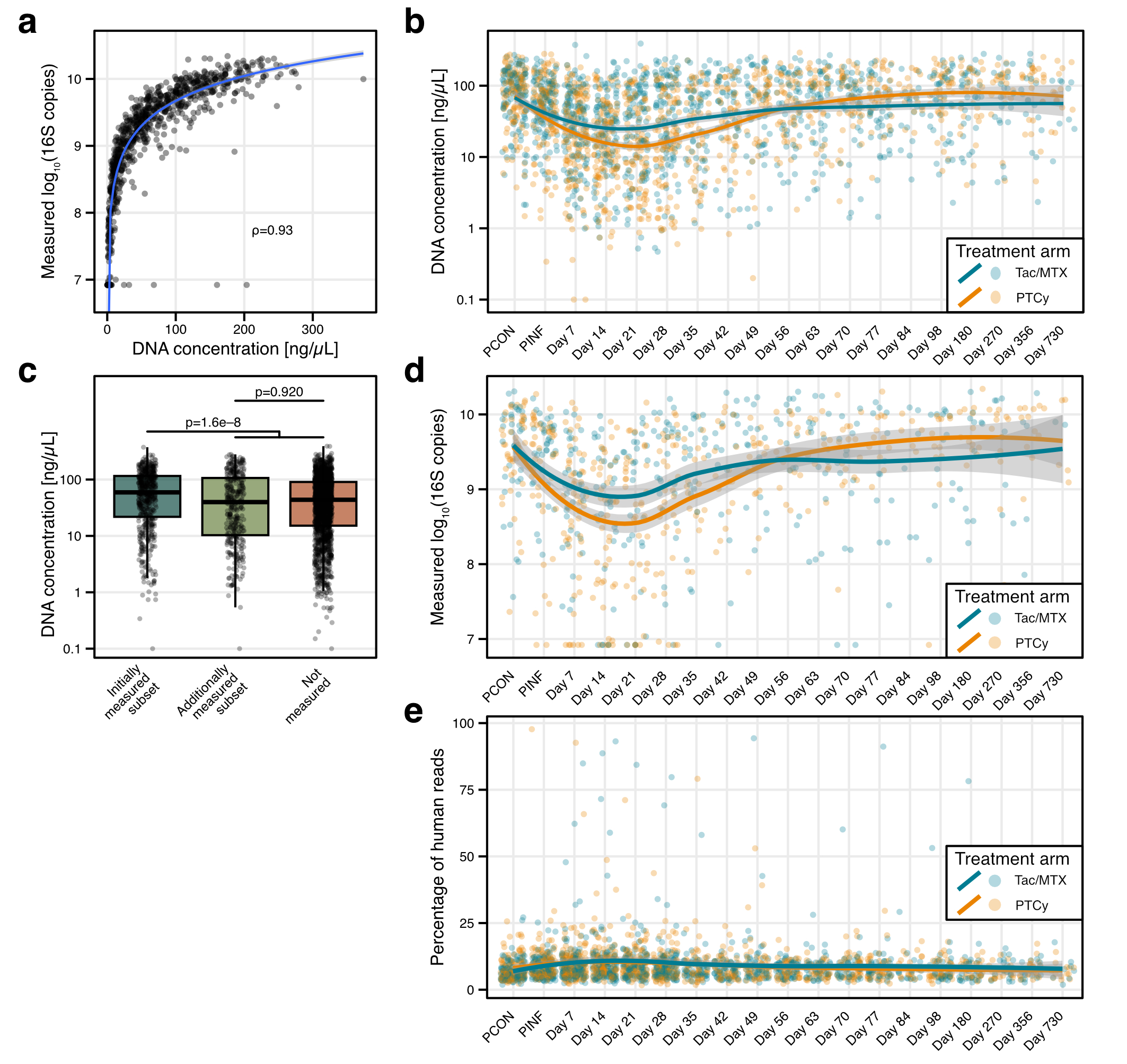

**SFig. 2: Relationship between DNA concentration and 16S rRNA copy number per extraction**

**a)** Scatter plot showing the correlation between total DNA concentration and the measured log10 16S copy number per extraction. The blue line and grey area correspond to a trendline fitted using a linear model with the formula y~log(x) and the 95% confidence interval. **b)** Total DNA concentration over concentration in both study arms. The trendline for each study arm was estimated using a LOESS model with the formula y~x, with the grey area showing the 95% confidence interval. PCON: pre-conditioning, PINF: pre-infusion. **c)** Distribution of DNA concentration for the first batch of samples measured via ddPCR, showing that the first batch consisted of samples with higher concentration, whereas the second batch of samples better reflected the overall distribution of DNA concentration. Differences in distributions were tested with a t-test. **d)** Log10-transformed 16S copy number per extraction over time, showing only samples that were measured via ddPCR. The trendline for each study arm was estimated using a LOESS model with the formula y~x, with the grey area showing the 95% confidence interval. **e)** Percentage of human reads within each sample over time, for both study arms. The trendline for each study arm was estimated using a LOESS model with the formula y~x, with the grey area showing the 95% confidence interval.

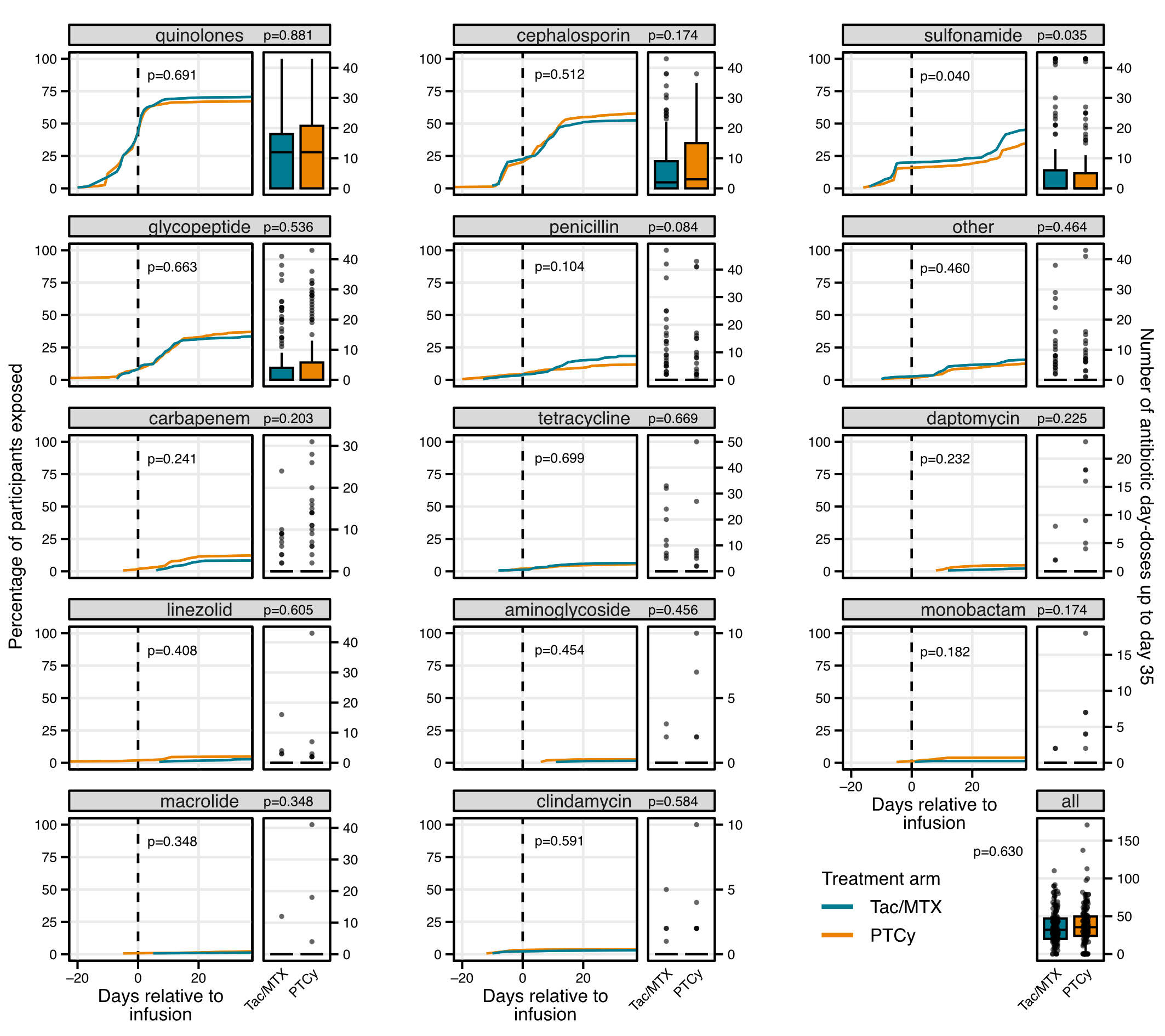

**SFig. 3: Exposure to different antibiotics classes up to day 35 post-HCT**

For each antibiotic class, the cumulative percentage of participants exposed to this antimicrobial class up to day 35 post-HCT is shown, split by treatment arm. Difference between distributions was calculated with a log-rank test. On the right hand side of each plot, the number of doses per participant are shown as box plots (see Figure 1 for box plot definitions). Difference between doses was tested with a Wilcoxon rank sum test.

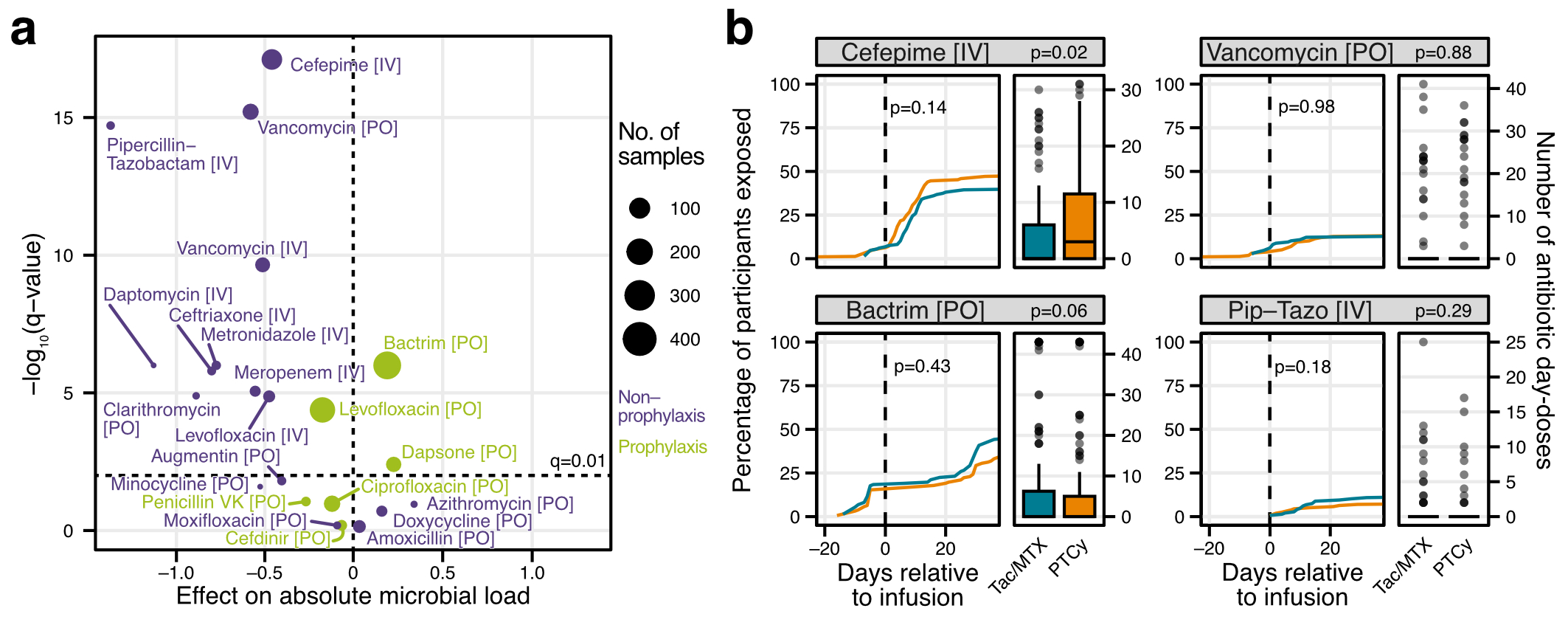

**SFig. 4: Effect of prophylaxis and non-prophylaxis antibiotics on total microbial load**

**a)** Volcano plot showing the significance and effect size of individual antibiotics on absolute microbial load in a sample, as estimated by a linear model. Each antibiotic (with its route of administration) was classified as either common prophylaxis or as non-prophylaxis (treatment for acute infections). Note that these classifications are broad and might differ from case to case, especially given differences in management and prophylaxis plans across participating medical centers. **b)** The cumulative percentage of participants exposed to different antibiotics up to day 35 post-HCT is shown, split by treatment arm, for selected antibiotics. Difference between distributions was calculated with a log-rank test. On the right hand side of each plot, the number of day-doses per participant are shown as box plots (see Figure 1 for box plot definitions). Difference between doses was tested with a Wilcoxon rank sum test.

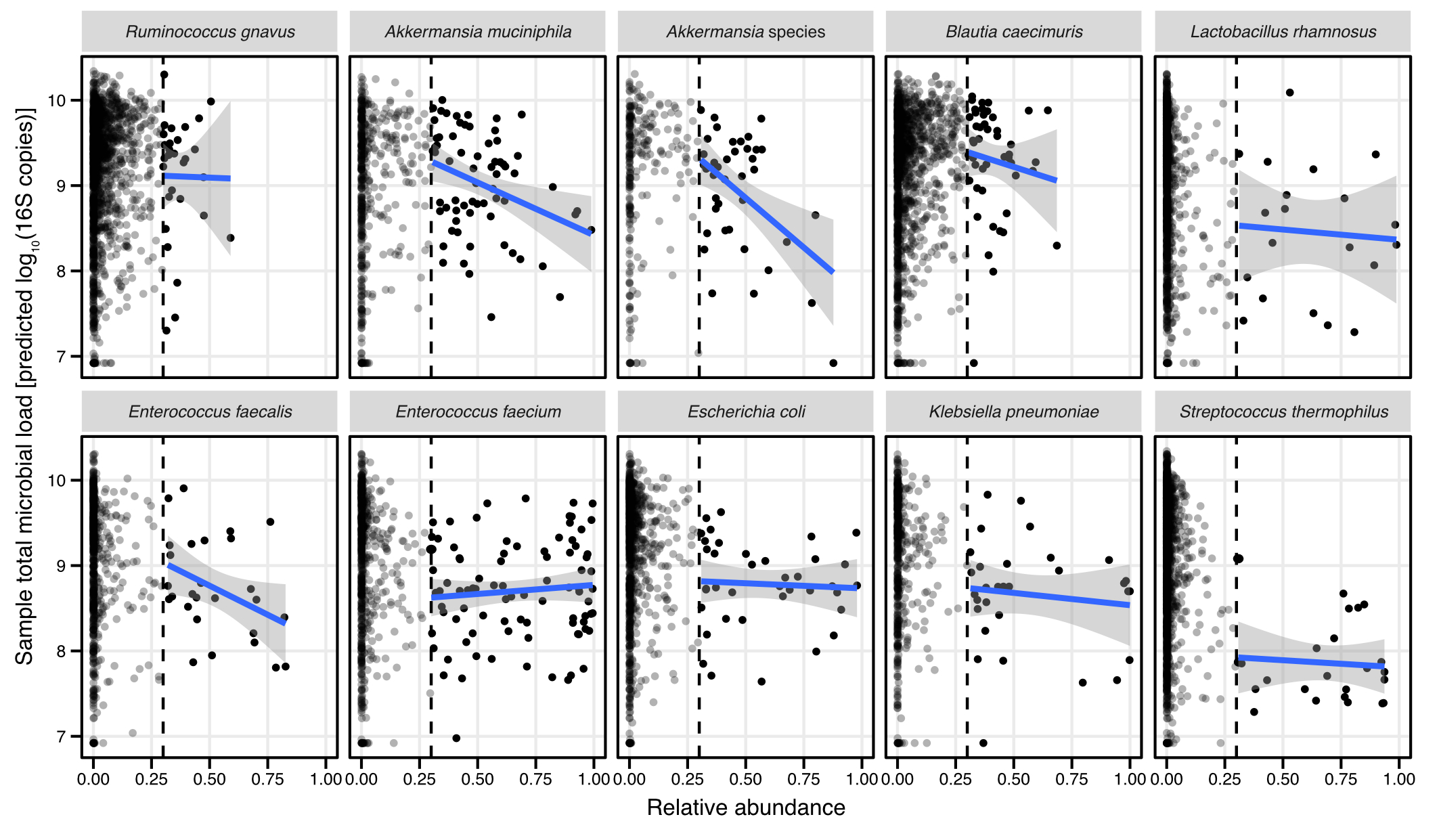

**SFig. 5: Relationship between relative abundance and total microbial load for commonly dominating species.**
For each of the most commonly dominating species, the relative abundance of the species in question is plotted against the log-transformed 16S copy number per extraction as proxy for the total microbial load per sample. In each panel, the data from all samples is included. For dominated samples (more than 30% relative abundance, see dashed black line), a trendline was fitted using a linear model with the formula y~x, shown as blue line, with the 95% confidence interval as grey area.

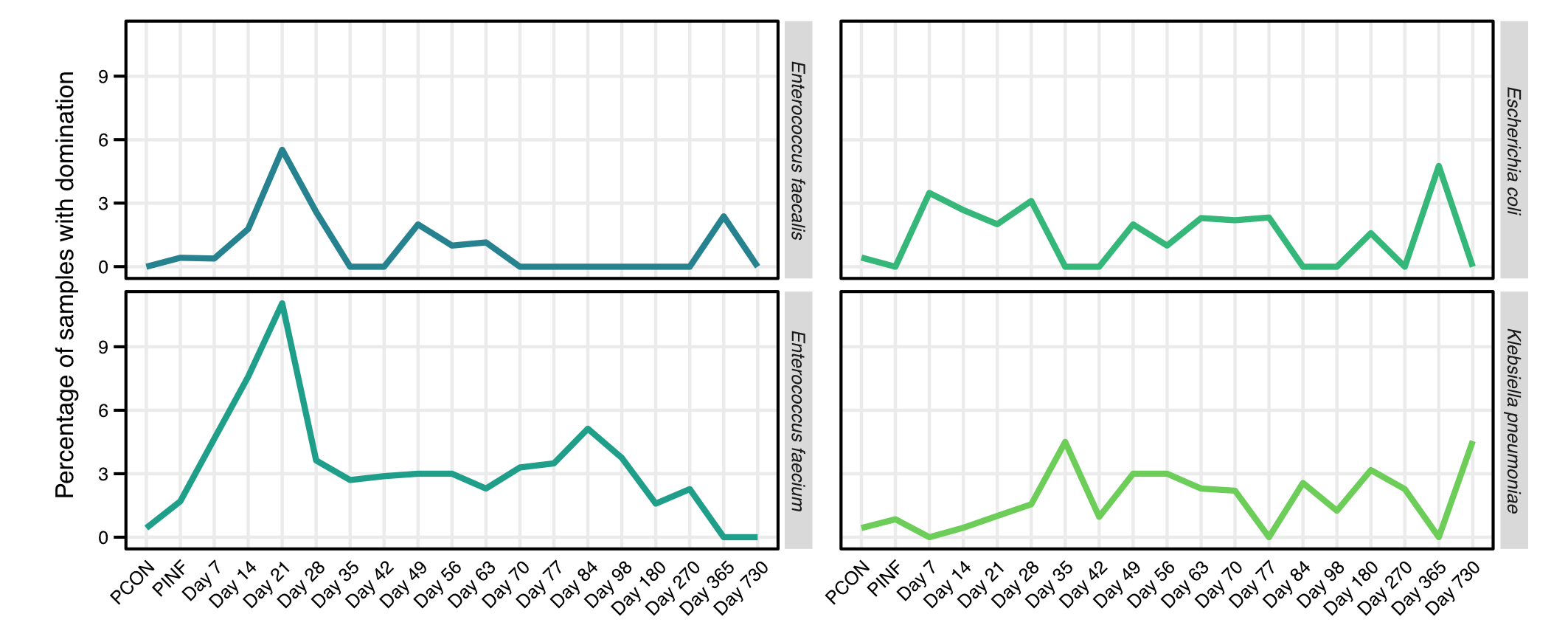

**SFig. 6: Domination rate for enteric pathogens over time**

Percentage of samples dominated (more than 30% relative abundance explained by a single species) shown as lineplots over the course of the study, for *E. coli*, *K. pneumoniae*, *E. faecalis*, or *E. faecium*.

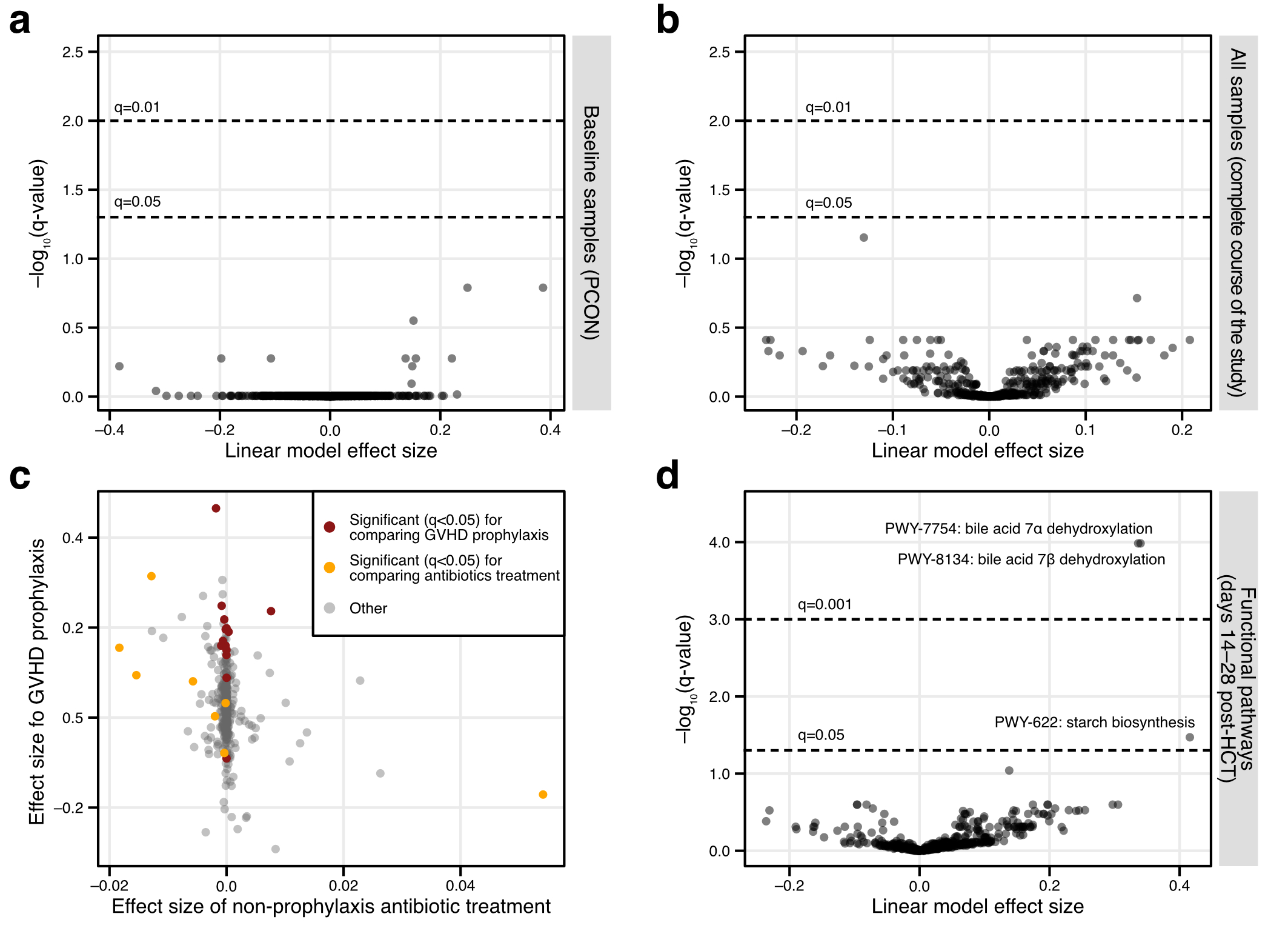

**SFig. 7: Differential abundance between study arms at baseline and over the complete study period**

**a)** Volcano plot showing the results of a linear model, comparing the relative abundance of individual microbial species across treatment arms for all baseline samples. **b)** Volcano plot showing the results of a linear model, comparing the relative abundance of individual microbial species across treatment arms for all samples donated over the course of the study. **c)** Linear model effect size for the model estimating the effect of non-prophylaxis antibiotics on individual microbial species between days 14–28 post-HCT plotted against the effect size for GVHD prophylaxis. Species with a q-value (p-value after correction for multiple hypothesis testing) less than 0.05 in either model are coloured. This plot indicates that species that show a strong association with GVHD prophylaxis are not confounded by antibacterial treatment. **d)** Volcano plot showing the results of a linear model, comparing the relative abundance of functional pathways across treatment arms for samples donated between day 14–28 post-HCT.

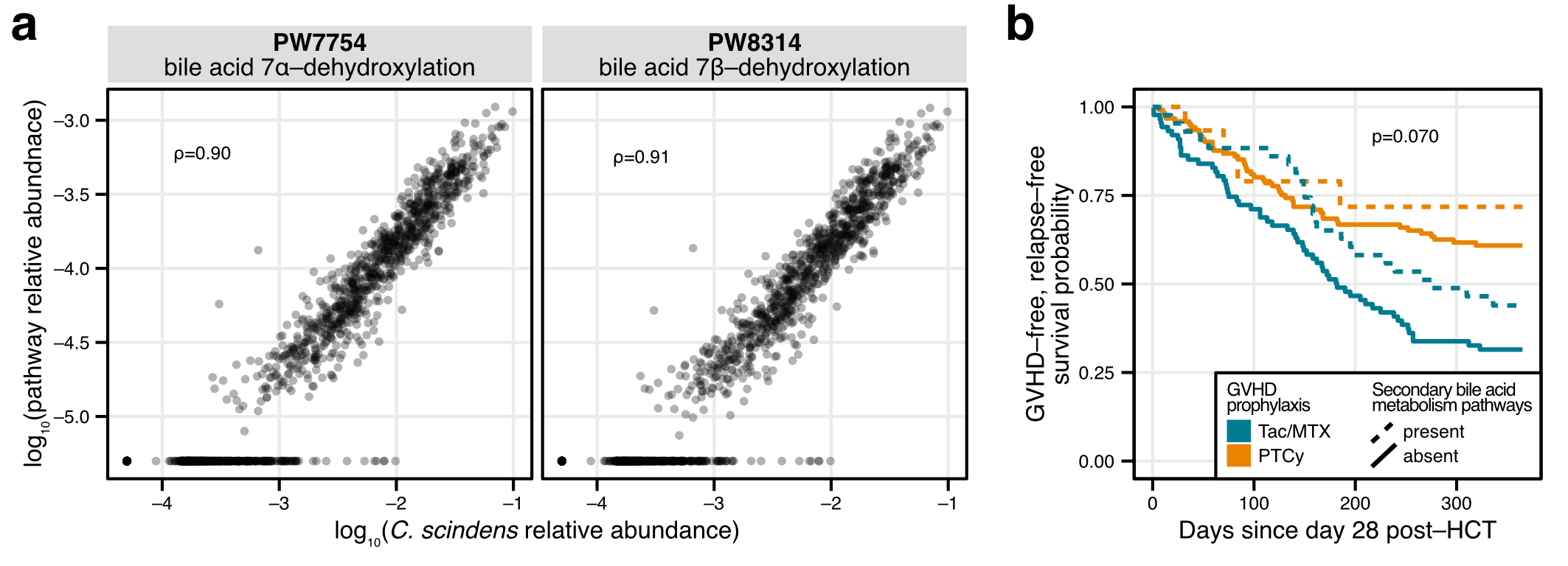

**SFig. 8: Correspondence between bile acid metabolism and *C. scindens* abundance**

**a)** Scatter plot showing the correspondence between the relative abundance of *C. scindens* and pathways involved in secondary bile acid metabolism. **b)** Kaplan-Meier probability curves for GVHD-free, relapse-free survival (GRFS), separated by GVHD prophylaxis and presence or absence of secondary bile acid metabolism pathways at days 14–28 post-HCT. P-value for the influence of secondary bile acid metabolism pathways was generated with a Cox regression model.

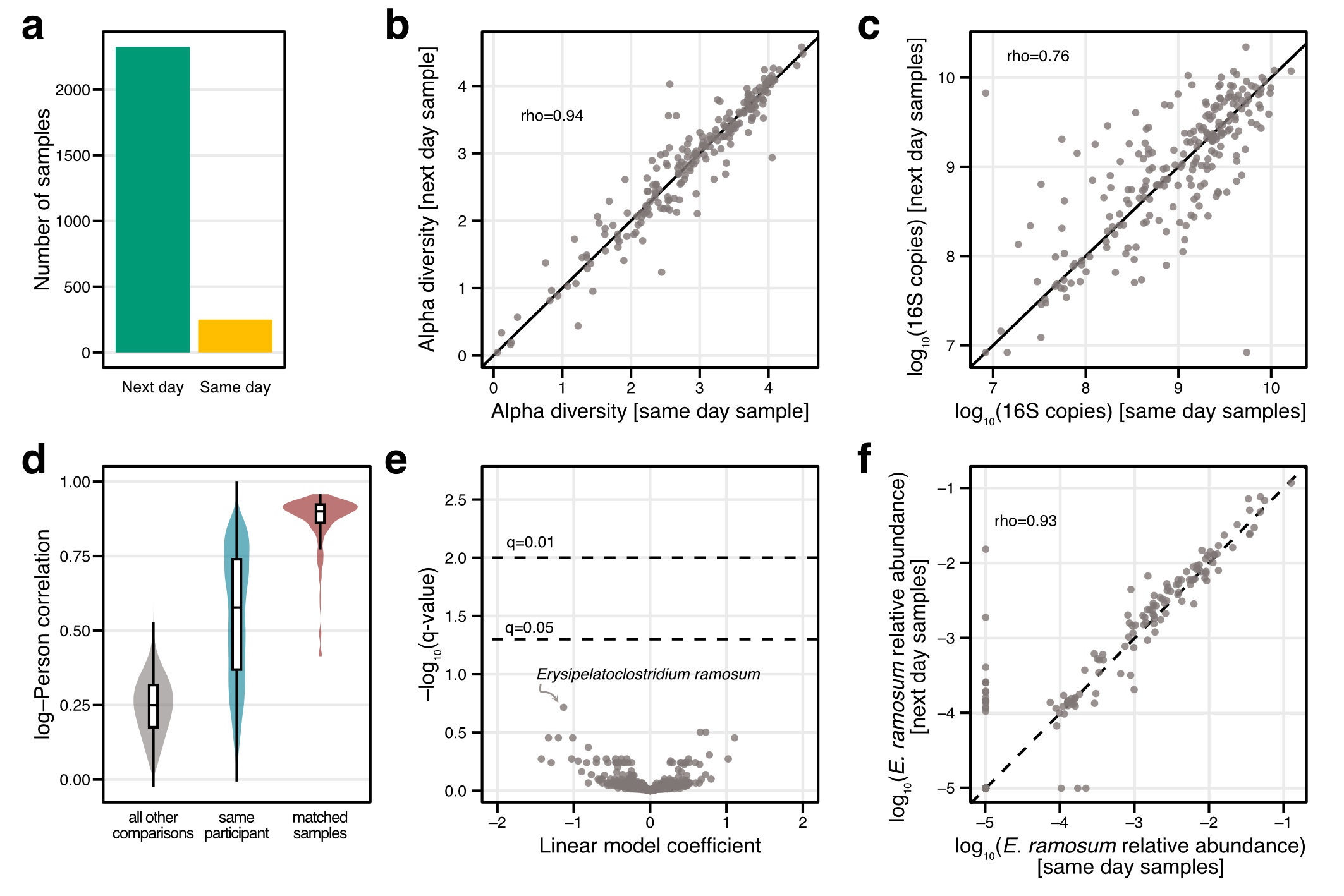

**SFig. 9: Comparison between same-day and next-day sample processing**

**a)** Number of samples collected with next-day or same-day processing. **b)** Alpha diversity (calculated using Shannon’s index) for samples processed using both next-day and same-day processing. Spearman correlation between alpha diversity values is 0.94, indicating high correspondence between processing techniques. **c)** Log10-transformed 16S copies for samples processed using both next-day and same-day processing. Spearman correlation is 0.76, indicating general concordance, yet some variability in 16S copies per extraction for processing techniques. Note that some values are the result of digital droplet PCR measurements and some values were predicted using our machine learning model. **d)** Pearson correlation values (calculated on log10-transformed relative abundance values) compared for matched samples (processed using both next-day and same-day processing), for samples from the same participant, and for unrelated samples, shown as violin and box plots. See Figure 1 for box plot definitions. **e)** Volcano plot showing the results of a linear model, comparing the relative abundance of individual microbial species for samples processed using both next-day and same-day processing. No significant differences after correction for multiple hypothesis testing were found. The most significant species is annotated. **f)** Scatter plot showing the relative abundance for *E. ramosum* (see e)) for samples processed using same-day and next-day sample processing. The spearman correlation is 0.93, indicating high concordance between processing techniques.
